# Comprehensive single cell profiling of ageing glial cells reveals impaired Wnt signalling and Jun transcription factors regulating cortical astrocytes

**DOI:** 10.64898/2026.04.16.718928

**Authors:** Maroussia Hennes, Manja Thorwirth, Chu Lan Lao, Jeff A. Stogsdill, Paola Arlotta, Judith Fischer-Sternjak, Maria L. Richter, Magdalena Götz

## Abstract

Understanding age-related cellular dysfunction in the brain is essential for developing strategies to promote healthy ageing. Towards this aim, we took advantage of a previously established mild dissociation method to profile cells in the cerebral cortex grey matter of adult and aged mice. This revealed glial cells with largely up-regulated and other glia and neurons with largely down-regulated gene expression upon ageing. Astrocytes were involved in increased interactions with microglia and decreased interaction with neurons, highlighting potent age-induced changes in their regulatory roles. Single cell RNA-seq and single nuclei multiome analysis of astrocytes uncovered down-regulation of Wnt-signalling with increased expression of its inhibitors and reduced RNA and protein levels of its effectors JunB/D, acting downstream of Wnt signalling in ageing. This was confirmed by RNA-scope and immunostainings, as well as in human data. Notably, injection of JunD-expressing viral vectors in astrocytes increased their proliferation and HMGB1 levels in the aged brain, indicative of a more youthful astrocyte state.

**Main points:**

- Transcriptomic analysis uncovers cell type–specific impact of ageing in the cortical grey matter, including altered intercellular communication networks.
- Multiomic profiling identifies dysregulated Wnt signalling in ageing cortical astrocytes.
- Ageing astrocytes exhibit upregulation of the Wnt signalling regulators Maml2 and Daam2, accompanied by downregulation of the AP-1 transcriptional complex component JunD.
- Overexpression of *JunD* increases proliferation after mild injury in aged astrocytes.

## Introduction

### Astrocytes as targets to comprehend and combat physiological brain ageing

Population ageing has become a defining demographic shift in modern societies, reflected by the continuously growing proportion of aged individuals. This rise in aged individuals goes hand in hand with an increase in age-related conditions including cognitive decline and neurodegeneration (World Health Organization, 2021; Fu et al., 2025; Peng et al., 2025). Advancing our understanding of brain ageing is imperative to unravel novel strategies capable of promoting healthy brain ageing.

Central nervous system (CNS) ageing is a complex, multicellular process that leads to progressive cellular deterioration, affecting the physiology of both neurons and glial cells such as astrocytes (López-Otín *et al*., 2023; Tartiere, Freije and López-Otín, 2024). Interestingly, from a transcriptomic point-of-view, astrocytes have been shown to be among the first cell types to be impacted by ageing (Ximerakis *et al*., 2019; Allen *et al*., 2023; Hahn *et al*., 2023; Jin *et al*., 2025). In the CNS, astrocytes are indispensable to ensure neuronal survival and maintain adequate brain function by mediating a variety of important processes, such as metabolic, antioxidant and synaptic activity (Barres, 2008; Allen, 2014; McBean, 2018; Verkhratsky *et al*., 2021, 2023). Moreover, astrocytes are recognized as a diverse cell population with distinct subtypes existing between and within brain regions, defined by molecular, morphological and functional differences (Ohlig *et al*., 2021; Bocchi *et al*., 2025; Hennes *et al*., 2025).

Previous research already indicated that both physiological and pathological ageing impacts astrocyte function in a region-specific pattern (Boisvert *et al*., 2018; Hahn *et al*., 2023). Transcriptomic analyses of aged brains indicate a marked shift toward astrocytic inflammatory phenotypes, evidenced by widespread upregulation of gliosis-associated transcripts predominantly in the white matter, such as Gfap and C4b, while simultaneously revealing ageing-related, region-specific alterations in astrocytic genes involved in synaptic regulation, including Sparc (Boisvert *et al*., 2018; Clarke *et al*., 2018; Ximerakis *et al*., 2019; Allen *et al*., 2023; Hahn *et al*., 2023; Linker *et al*., 2025). Ageing also triggers the emergence of astrocyte substates detrimental to brain function (Lee *et al*., 2022; Linker *et al*., 2025). Lee et al, for instance, revealed the emergence of autophagy-dysregulated astrocytes in the ageing hippocampus, affecting their capacity to regulate synaptic homeostasis (Lee *et al*., 2022). Ageing-related impairments in astrocytic synaptic functions may also arise from their reduced morphological complexity in the aged brain, limiting the ability of distal processes to effectively cover and modulate nearby synapses (Popov *et al*., 2021, 2023).

Astrocytes evidently play a significant role in brain ageing, yet it is less recognized that their ageing phenotype may initiate neuronal impairment. Several neurological conditions have already been re-classified as primary gliopathies, with glial dysfunction implicated not only in disease progression but also onset (Verkhratsky *et al*., 2012, 2023). Increasing evidence that aged astrocytes can drive early neuronal deficits suggests that age-related cognitive decline and brain disease may likewise be gliopathic in origin, with neurodegeneration arising as a downstream consequence. Interestingly, it has recently been shown that astrocytes can switch between cellular ‘substates’, meaning that the same astrocyte can initially be neuroprotective before becoming neurotoxic (Zhang *et al*., 2025). This raises the possibility that ageing-related astrocyte states are not fixed and could be shifted toward a more youthful phenotype, offering new avenues for therapeutic intervention.

While next-generation sequencing (NGS) has greatly advanced our understanding of brain physiology in health and disease, discrepancies across studies highlight the need for integrated multi-omics approaches optimized from sample preparation through analysis. In this project, we aimed to further characterize the ageing grey matter (GM) of the cerebral cortex using complementary omics approaches and protocols optimized for aged tissue. First, we assessed the global impact of ageing across all identified cell types, including associated changes in intercellular communication, to generate a comprehensive overview of those most affected cell types and clusters, based on both quantitative and qualitative transcriptomic alterations. We then focused specifically on astrocytes, as multiple of their interactions with other cell types were changed in physiological ageing. In addition to the prevailing focus on astrocytic inflammatory and reactive pathways increased in ageing, our integrated multi-omics profiling consistently detected altered Wnt signalling. Interestingly, these ageing-induced alterations in Wnt signalling are seemingly cluster-specific and can be validated across datasets and species. Importantly, we further identified members of the AP-1 transcriptional complex, acting downstream of Wnt signalling to be reduced in aged cortical astrocytes and, when overexpressed, restoring proliferative capacity of mildly reactive astrocytes. This is indicative of a more youthful astrocyte state, as proliferation of reactive astrocytes is strongly decreased in ageing (Heimann *et al*., 2017). Thus, our work identified a key pathway regulating astrocyte ageing and demonstrated that its effects can be partially reversed.

## Material and Methods

### Experimental animals

C57BL6/J mice (Charles River Laboratories), Aldh1l1-eGFP mice (Tg(Aldh1l1-EGFP)OFC789G-sat/Mmucd, Gensat Project) (Heintz, 2004) and Tcerg1l-CreERT2/tdTomato mice (Matho *et al*., 2021); Jackson stock # 007914) were used for this study. Adult mice were 3-4 months old and aged mice were used between 22-24m of age. All animals were housed under standard laboratory conditions under a 12h/12h light/dark cycle with food (Altromin, 1310M) and water available *ad libitum* at the Core Facility Animal Models, Biomedical Center, Faculty of Medicine, LMU Munich under specific-pathogen-free conditions. Experimental procedures were performed in accordance with animal welfare policies and were approved by the Government of Upper Bavaria (Germany).

### Single nuclei isolation and sequencing

The protocol was adapted from Thrupp et al., 2020 (Thrupp *et al*., 2020). Briefly, nuclei were isolated from freshly dissected adult (n = 1) and aged (n = 1) mouse cortices in ice-cold HBSS (Gibco) supplemented with 1% HEPES (Gibco) solution. Cortices were sliced and manually homogenized using a pestle in 1mL of homogenization buffer [HB; Sucrose (320mM), Calcium chloride (5mM), Magnesium Acetate (3mM), Tris (10mM), Ethylenediaminetetraacetic acid (0.1mM), Igepal (0.1%), Phenylmethylsulfonylfluoride (0.1mM), 2- Mercaptoethanol (1mM), MQ water) with 5µl RNasin Plus]. The homogenate was strained using a 70µm strainer and washed with 1.65mL HB to a final volume of 2.65mL. An equal volume of Gradient medium (Calcium chloride (5mM), Optiprep (50%), Magnesium Acetate (3mM), Tris (10mM), Phenylmethylsulfonylfluoride (0.1mM), 2-Mercaptoethanol (1mM) was added (Vf = 5.3mL). To isolate the nuclei, first an Optiprep Diluent medium (ODM; Potassium chloride (150mM), Magnesium chloride (30mM), Tris (60mM), Sucrose (250mM) was prepared. The sample was added to a 4mL 29% cushion (Optiprep (29%), ODM). The sample was centrifuged at 10 000g for 30 minutes at 4°C. The supernatant was removed and nuclei were resuspended in 200mL resuspension buffer (10xPBS (1x), BSA (1%), RNasin Plus (0.2U ml −1)). Clumps were disrupted by pipetting and filtered through a 70 followed by a 35µm cell strainer.

Single nuclei suspensions were processed using the 10X Genomics Chromium Next GEM Single Cell Multiome ATAC + Gene Expression Reagent Kit according to manufacturer’s instructions. Illumina libraries were sequenced on the NovaSeq 6000 platform after quality assessment with the Bioanalyzer (Agilent Technologies), with an average read depth of 25 000 raw reads per cell.

### Single cell isolation and sequencing

Single cell samples from adult cortices (n = 3) were reused from Bocchi et al (Bocchi *et al*., 2025). Aged single cell suspensions from mouse cortices (n = 2) were prepared as previously described using the Papain Dissociation System (Worthington Biochemical) followed by the Dead cell Removal kit (cat. No. 130-090-101, Miltenyi Biotec) (Bocchi *et al*., 2025). Cells were suspended in 1xPBS with 0.04% BSA for a final concentration of 1000 cells/µl. Single cells were further processed using the Single Cell 3’Reagent Kits v3.1 from 10x Genomics. After quality assessment with Bioanalyzer (Agilent Technologies), Illumina libraries were sequenced on the NovaSeq 6000 platform with an average depth of 30 000 raw reads per cell.

For single cell samples to isolate astrocytes from different layers 300ffim thick vibratome sections were prepared from cerebral cortices of Tcerg1l-CreERT2 (Matho *et al*., 2021) crossed with the ROSA26loxSTOPlox-tdTomato reporter allele obtained from Jackson Laboratories (stock # 007914) to obtain Tcerg1l-CreERT2/tdTomato mice (Stogsdill *et al*., 2022), labelling layer 5 neurons in S1 to allow manual dissections of the cortex layers: first layer 1 was removed, then the layers above the fluorescently labelled layer 5 was cut out as “Layer2-4”, followed by collecting the fluorescent “layer5” and the remaining band above the WM as “Layer6”. These samples were pooled, dissociated and astrocytes were isolated by MACS using the ACSA2 antibody as described in Ohlig et al., 2021 without myelin removal kit (Ohlig *et al*., 2021). Ependymal cell contamination as discovered by Ohlig et al. 2021 was not a problem here, as WM and the underlying ependymal layer were removed during dissection (Ohlig *et al*., 2021). For sequencing, cells were processed further as described in Stogsdill et al. 2022 (Stogsdill *et al*., 2022) using the Chromium Single Cell 3’ Kit v3.1 (10x Genomics, PN-1000121).

### Analysis of 10X genomics scRNA-seq of adult and aged cortical GM samples

For better comparability of the adult and aged cortical GM samples, data were processed as described in Bocchi et al (Bocchi *et al*., 2025). In brief, reads were aligned to mm10 using *CellRanger*-7.0.1. To remove ambient RNA, *CellBender* version 0.3.0 was applied on the raw count matrix. Genes were kept if they were detected in a minimum of 3 cells. Cells were filtered to have between 350 and 5000 genes and less than 15% mitochondrial reads. Count normalization was done using *SCTransform* and replicates were integrated using *Harmony*. Cell type annotation was done based on *leiden* clustering using classical marker genes from literature, e.g. *Sox9*, *Gfap*, *Aldh1l1*, *Slc1a2*, *Slc1a3*, *S100b*, *Gja1*, and *Fgfr3* for astrocytes. Differential expression analysis between aged and adult was performed per cell type using *scanpy’s sc.tl.rank_genes_groups*. Differential cell-cell-communication was assessed using *Community* (Solovey *et al*., 2024). Clustering of astrocytes was done at a resolution of 0.25, resulting in 5 clusters.

### Analysis of 10X genomics scRNA-seq of layer-separated data

Reads were aligned to mm39 using *CellRanger*-9.0.1. To remove ambient RNA, *CellBender* version 0.3.0 was applied on the raw count matrix. Cells were kept if they had a minimum of 500 genes. Genes were kept if they were present in at least 3 cells. Finally, cells with more than 25% mitochondrial reads were removed. Library size normalization was done using *scran* (L. Lun, Bach and Marioni, 2016). Cell type annotation was done based on *leiden* clustering using classical marker genes from literature, e.g. *Sox9*, *Gfap*, *Aldh1l1*, *Slc1a2*, *Slc1a3*, *S100b*, *Gja1*, and *Fgfr3* for astrocytes. Differential expression analysis between layers was done using *scanpy’s sc.tl.rank_genes_groups (Wolf, Angerer and Theis, 2018)*. The top 12 most differentially expressed genes per layer were selected with adjusted p-value < 0.05, presence in at least 5 cells, and log2 fold change > 1.

### Analysis of human single nuclei data

For cross-species validation the publicly available human snRNA-seq dataset from Chien et al was used (Chien *et al*., 2024). This dataset contains snRNA-seq data from data healthy young (23-30 years old) and aged (70-74 years old) human prefrontal cortex (Chien *et al*., 2024). The processed data was downloaded from: https://cellxgene.cziscience.com/collections/91c8e321-566f-4f9d-b89e-3a164be654d5.

### Analysis of multiome RNA and ATAC

Multiome data of combined RNA and ATAC were aligned to mm10 using C*ellRanger-arc* version 2.0.1. For the ATAC part, fragment files were combined for the two age conditions and peaks were called using *macs3*. The *muon* package was used to load the data into python. For the RNA part, genes were kept if they were present in at least 5 cells. Cells were kept if they had between 1000 and 7000 genes, a maximum of 50,000 reads, and a maximum of 5% mitochondrial reads. Doublets were removed using *scrublet* with a threshold of 0.1. Library size normalization was done with *sc.pp.normalize_total* with a target sum of 10,000 reads. Cell type annotation was done based on *leiden* clusters using classical cell type marker genes from literature (same as above). Differential expression analysis between aged and adult was performed per cell type using *scanpy’s sc.tl.rank_genes_groups*. *Episcanpy’s epi.tl.find_genes* was used to associate peaks to genes that are within 2kb downstream of the peak (Danese *et al*., 2021).

For a combined multiomic analysis, SCENIC+ was used (González-Blas *et al*., 2023). Based on the model outputs, 60 topics were selected for the cistopic part of the analysis. Astrocytes were selected and eRegulon specificity was calculated for the two age conditions to get agespecific eRegulons in astrocytes.

### Tissue preparation, RNAscope and immunohistochemistry

Mice were perfused transcardially under anesthesia [ketamine (100 mg kg^−1^)/xylazine (10 mg kg^−1^)] with 1x phosphate-buffered saline (PBS) followed by 4% paraformaldehyde (PFA). Postfixation was performed overnight (ON) with 4% PFA at 4°C. After 1x PBS wash, brains intended for RNAscope were transferred to a 30% sucrose solution in PBS. Coronal brain sections of 30µm were prepared using a cryostat. For immunofluorescence analysis, 50µm thick coronal sections were cut using a Leica VT1000S vibratome.

RNAscope was performed using the RNAscope Multiplex Fluorescent Reagent kit v2 (ACD-Bio) according to the manufacturer’s instructions. For hybridization, the following probes were obtained from ACDBio: *Daam2* (probe-Mm-*Daam2*-C1 Manual assay), *Maml2* (Probe-Mm-*Maml2*-C1 Manual assay), *Tcf7l2* (probe-Mm-*Tcf7l2*-C1 Manual assay), *Znrf3* (probe-Mm-*Znrf3*-C3 Manual assay) and *Slc1a3* (probe-mM-*Slc1a3*-C2 Manual assay).

For immunostaining, a heat-induced epitope retrieval (HIER) was performed at 80°C for 30min in a 10mM sodium citrate buffer (pH=6) on free-floating sections. Next, sections were incubated for 45min in blocking solution (PBS with 5% BSA and 0.5% Triton X-100).

The following primary antibodies were used and incubated ON at 4 °C in antibody solution (PBS with 2%BSA and 0.5% Triton X-100): rabbit anti-JunD (1:500 dilution, Abcam 181615); rabbit anti-cJun (1:500 dilution, Cell Signalling 9165T); rabbit anti-JunB (1:200 dilution, Cell Signalling 3753); mouse anti-S100b (1:500 dilution, Sigma S2532); chicken anti-RFP (1:500 dilution, Rockland 600-901-379S); chicken anti-GFP (1:300 dilution; Aves Labs GFP-1020); goat anti-Sox9 (1:1000 dilution, R&D Systems AF3075-SP); rabbit anti-Hmgb1 (1:1000 dilution, Abcam ab18256). After washing (1x PBS, 3× 10 min at RT), secondary antibodies were incubated at room temperature for 2h: anti-rabbit 647 (1:1000, Thermo Fisher Scientific); anti-mouse 488 (1:1000, Thermo Fisher Scientific); anti-chicken 488 (1:1000, Thermo Fisher Scientific); anti-chicken 594 (1:250, Thermo Fisher Scientific); anti-goat 488 (1:250, Thermo Fisher Scientific). For nuclear staining: sections were incubated with 4’,6-diamidino-2-phenylindole (DAPI; 0.1 mg/ml cat.no. D9564-10 mg, Sigma-Aldrich) together with the secondary antibodies. 5-EdU incorporation was visualized with the ClickiT EdU Alexa Fluor 647 Imaging kit (Thermo Fisher Scientific).

### Image acquisition, processing and analysis

For immunostaining, confocal microscopy was performed using a Zeiss LSM710 microscope with ZEN software (Black edition, v.2.3 SP1, Zeiss) or a Leica Thunder Imager (V.3.5.7.23225, Leica). Per animal, 2-3 images were acquired with a x25/0.80 or a x40/0.95 and analysed using ImageJ (V.2.14.0/1.54f, NIH). In each section, cortical bins were defined, and the number of positive cells was quantified across all z-planes of the optical stack.

For RNAscope analysis, confocal microscopy was performed using a Leica SP8 or Stellaris confocal microscope with LASX software (V.3.5.7.23225, Leica). A x40/1.3 or x40/1.1 objective was used to acquire 1-2 sections per animal. Positive cells were quantified as described above. Intensity analysis was performed by defining a region of interest (ROI) encompassing the nucleus and measuring the mean grey value. Background signal was estimated by placing an ROI of identical size in an adjacent area lacking specific signal. For subsequent statistical analyses, nuclear intensities were normalized to their corresponding background values.

For RNAscope analysis of *Maml2*, one section per animal was analysed using QuPath (Bankhead *et al*., 2017). Cell detection was performed using the ‘cell detection’ module using DAPI to detect the region of interest. Using the ‘subcellular detection’ module, the quantity of distinct punctuate dots was counted in each cell. A threshold of more than 3 estimated dots per cell for each candidate and more than 15 estimated dots per cell for *Slc1a3* was established.

Data analysis was performed blinded. All statistical tests were performed using Prism v10.4.1 (GraphPad Software) or R v4.5.2. Statistical significance was defined as *P<0.05, **P<0.01 and ***P<0.001. All column graphs are expressed as mean ± s.e.m. Normality assessment of the dataset was assessed using the Shapiro-Wilk test. Statistical differences between groups were analysed using unpaired t-tests for normally distributed data and Mann–Whitney tests for non-normally distributed data. To asses differences in layer distribution, Jun+JunD+JunB+/S100b+ cells were counted in bins representing Upper (0-30% section); Middle (30-70%) and Lower (70-100%) layers of the cortex.

### Lentiviral vector cloning and production

The plasmids pLV-CBh-mScarlet and pLV-CBh-3xFlag-mJunD-IRES-mScarlet were cloned via Gateway cloning. In brief, the destination plasmid pLV-CBh-DEST was derived from the pLenti7.3/V5-DEST™ Gateway™ Vector (Life Technologies), in which the CMV promoter was replaced with the hybrid CBh promoter (Addgene 50718). The mouse JunD (NM_010592.5) and the FLAG tag (DYKDHDG-DYKDHDI-DYKDDDDK) on the N-terminus were synthesized by Genewiz (Azenta) with a BglII/BamHI restriction enzyme site and subcloned into pENTR1A vector with IRES-mScarlet, mScarlet (Addgene 85042). For the MK-G (Mokola-G envelope)-pseudotyped lentivirus (LV), standard triple transfection of 293T cells (CRL-3216, ATCC) with an HIV-based, self-inactivating 2nd generation packaging plasmid pCMVΔR8.91 (kindly provided by Pavel Osten) and pseudotyping plasmid pMK-G (kindly provided by Philip Zoltick) was used. The LVs were concentrated from supernatants of transfected packaging cells by ultracentrifugation following standard protocols and resuspended in a buffer containing 50 mM Tris-HCl pH 7.8, 130 mM NaCl, 10 mM KCl, and 5 mM MgCl2.

### Stereotactic lentivirus injection

The MK-G pseudotyped lentiviral vectors, previously validated for astrocyte targeting (Natarajan *et al*., 2024), containing JunD cDNA and mScarlet reporter gene (pLV-Cbh-3xFLAGmJunD-IRES-mScarlet) were injected into the grey matter cortex of aged C57BL/6J mice. Hereto, mice were given an intraperitoneal injection containing fentanyl (0.05 mg/kg, Janssen), midazolam (5 mg/kg, Roche) and medetomidine (0.5 mg/kg, Fort Dodge). LV unilateral injections (titer: 1.93 × 109 transducing units (TU)/ml, 1 µl/injection side) were performed using coordinates relative to the bregma with an automated nanoinjector (Nanoliter 2010, World Precisions Instruments) at a slow speed (40 nl/min): anteroposterior = −1.0; mediolateral = +2.0; dorsoventral = −0.5. Anesthesia was reversed via subcutaneous administration of atipamezole (2.5 mg/kg, Janssen), flumazenil (0.5 mg/kg, Hexal) and buprenorphine (0.1 mg/kg, Essex). 5-EdU (Thermo Fisher Scientific) was administered via drinking water at a concentration of 0.2 mg/ ml containing 1% sucrose for 7 days before sacrificing the mice.

### Data availability

All raw sequencing data have been deposited in BioStudies under accession number S-BSST2601 (https://www.ebi.ac.uk/biostudies/studies/S-BSST2601). Adult grey matter scRNA-seq samples were taken from Bocchi et al., 2025. The publicly available human snmCT data were downloaded from https://cellxgene.cziscience.com/collections/91c8e321-566f-4f9d-b89e-3a164be654d5.

### Code Availability

Scripts used for data analysis are deposited at https://github.com/PhysiologicalGenom-icBMC/AgingAstrocytes.git

## Results

### Single-cell transcriptomic analysis of cortical GM indicates that glial cells are most affected by ageing

To obtain an overview of how ageing affects different cortical cell types and their intercellular communication, we took advantage of our previously established dissociation method, allowing unbiased glial cell capture and analysis even in tissue that is difficult to dissociate (Ohlig *et al*., 2021; Bocchi *et al*., 2025; Hennes *et al*., 2025) to perform single cell RNA sequencing (scRNA-seq) analysis. Single cells were isolated from cortical GM of adult (3m-4m) and aged (1y10m-2y) C57BL/6J mice. Quality control filtered cells based on gene counts (>350 and <5000) and mitochondrial gene percentages (<15%; Extended Data Fig. 1A-B). After filtering, this dataset contained 20,332 cells and 20,746 genes (Fig. 1A). Leiden clustering identified 9 distinct cell types, including astrocytes (Fig. 1B). Annotation of the different cell clusters was accomplished using specific cellular markers (Fig. 1B; Extended Data Fig. 1C-D).

**Figure 1.**
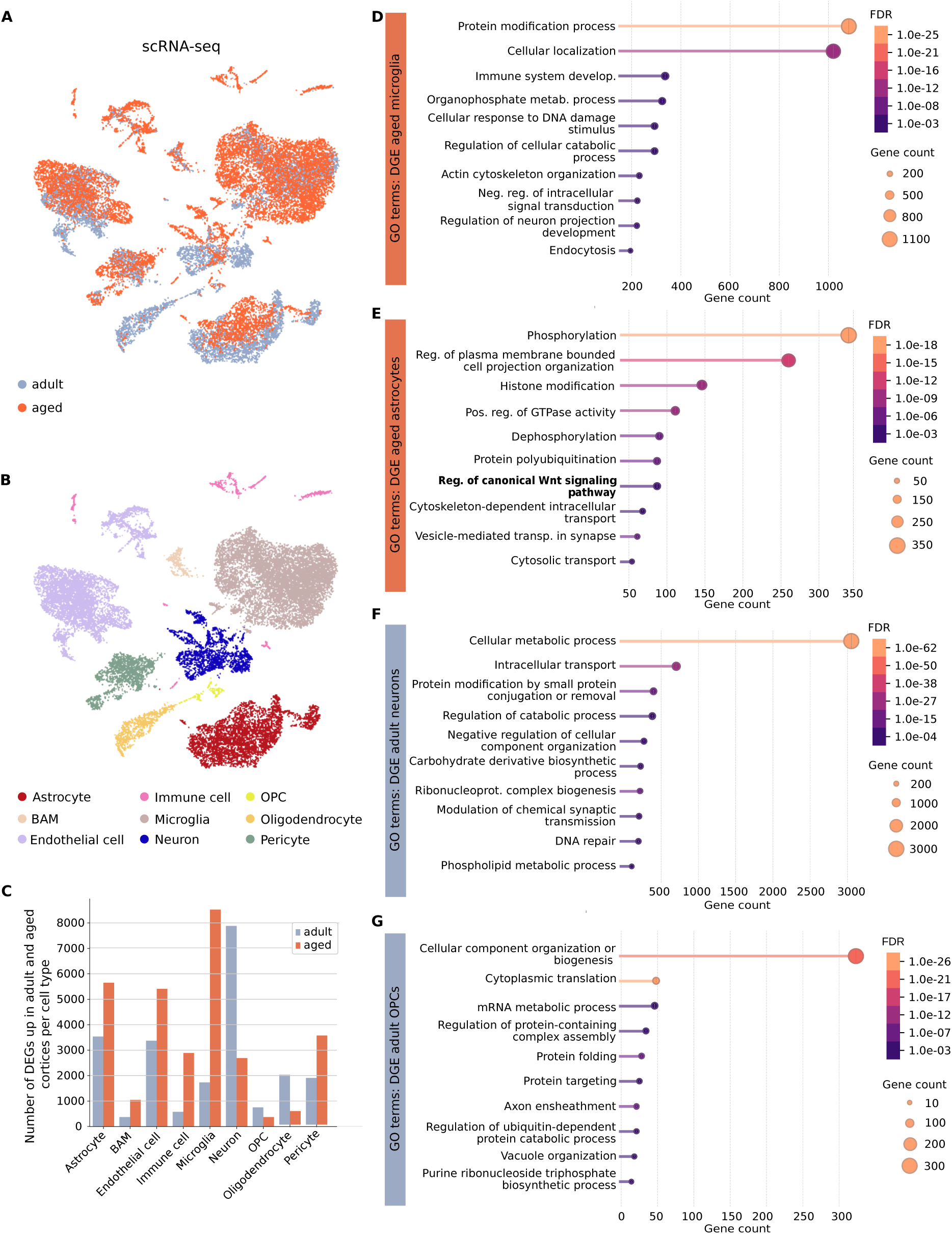
Single cell RNA-seq based profiling of adult and aged mouse cortical cell types. **(A)** Umap of scRNA-seq data from adult (colour-coded in grey, n = 3) and aged (colour-coded in orange, n = 2) mouse cortices. **(B)** Umap of cell cluster annotation, colour-coded by cell type. BAM = border-associated macrophage, OPC = oligodendrocyte precursor cell. **(C)** Bar plot representation of the number of significantly up-regulated genes per cell type and per condition (t-test, Benjamini-Hochberg adjusted p-value < 0.05). **(D)** STRING-based GO terms of genes significantly enriched in aged compared to adult microglia. **(E)** STRING-based GO terms of significantly enriched in aged compared to adult astrocytes. **(F)** STRING-based GO terms of genes significantly downregulated in aged compared to adult neurons. **(G)** STRING-based GO terms of genes significantly downregulated in aged compared to adult OPCs.

The number of differentially expressed genes in aged and adult cortices per cell type revealed a clear distinction between cell types showing predominantly age-related upregulation of genes, such as brain-associated macrophages (BAMs), immune cells, astrocytes and microglia, and those exhibiting mainly age-associated downregulation, including neurons and oligodendrocytes and their progenitors (OPCs) (Fig. 1C). In aged microglia, significantly upregulated genes are associated with Immune system development and Endocytosis (Fig. 1D), while in astrocytes they are related to Cell projection organization, Wnt signalling and Vesicle-mediated transport at the synapse (Fig 1E). In aged neurons, significantly downregulated genes are related to synaptic transmission and DNA repair (Fig. 1F), while in aged OPCs downregulated genes are involved in mRNA metabolic processes and regulation of ubiquitin-dependent protein catabolic processes (Fig. 1G).

Beyond up- or downregulation of genes, ageing also induces qualitative transcriptional changes that could undermine stable cell identity (Mertens *et al*., 2021; Yang *et al*., 2023; Gorelov and Hochedlinger, 2024). To visualize this, the gene expression profiles of all cell types in the adult and aged cortex (Extended Data Fig. 1E) were correlated with each other. This correlation matrix indicated that for some cell types such as endothelial and immune cells, the overall gene expression profile remained constant between adult and aged conditions. In contrast, microglia, astrocytes, oligodendrocyte progenitor cells (OPCs), and neurons exhibited the most divergent gene expression profiles between adult and aged states (Extended Data Fig. 1E). This prompted us to explore, how intercellular signalling between these cell types is altered in the aged condition.

### Ageing-induced changes in cell-cell interactions

A cell-cell interaction analysis was performed to assess how ageing affects the dynamics of intercellular communication. Figure 2A visualizes the cellular interactions ongoing in the adult and the aged cortex, illustrating major changes in signalling between cells in the aged condition. Increased cell-cell communication occurred mostly for microglia and immune cells, but also astrocytes increased their communication with microglia and immune cells in the aged condition, while they decreased their communication with neurons. These data are consistent with the concept that astrocytes may aggravate the neuroinflammatory conditions in ageing and reduce their support for neurons (Labarta-Bajo and Allen, 2025).

**Figure 2.**
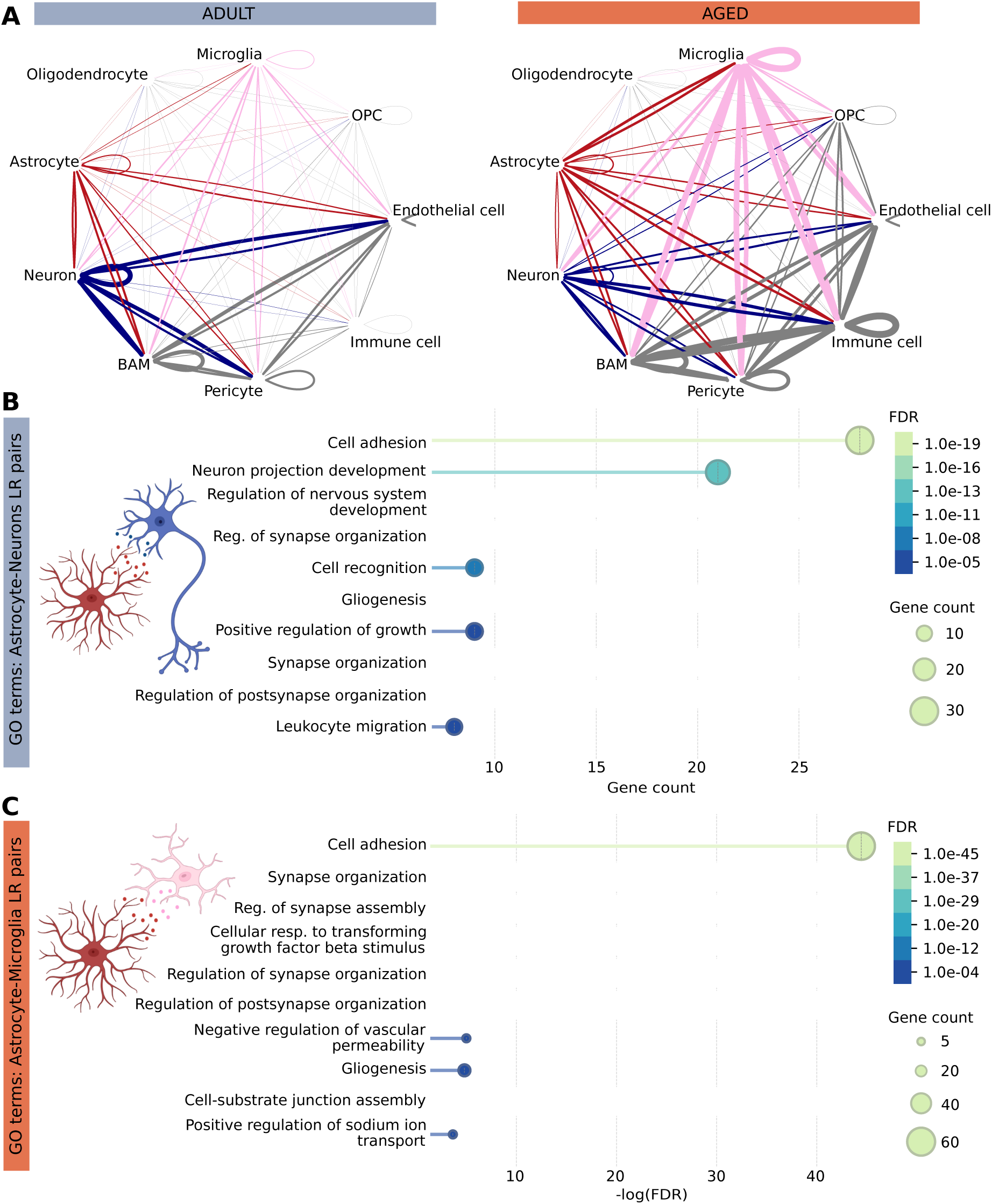
Ageing-induced alterations in cellular interactions in the mouse cortex. **(A)** Visualization of cell-cell interactions based on ligand-receptor pairs in adult and aged murine cortical cell types (red: involving astrocytes, pink: involving microglia, blue: involving neurons, grey: other). **(B)** STRING-based GO terms of age-associated ligand-receptor pairs involved in down-regulated astrocyte-neuron interactions. **(C)** STRING-based GO terms of age-associated ligand-receptor pairs involved in up-regulated astrocyte-microglia interactions. Illustrations of cell-cell-interactions were created in BioRender. Masserdotti, G. (2026) https://BioRender.com/rm6hq1j.

Ligand-receptor (LR) pairs underlying the downregulated astrocyte-neuron interactions pointed towards multiple GO terms associated with synapse organization (Fig. 2B). Among the downregulated astrocyte–neuron interactions, we identified the Brevican (Bcan)–Neuronal Cell Adhesion Molecule (Nrcam) ligand–receptor pair. Bcan has recently been identified as a marker of brain ageing, with its dysregulation affecting multiple cortical and subcortical structures (Liu *et al*., 2025). Further analysis of the genes associated with our identified synapse-related GO terms revealed a single gene common to all terms: *App*. Among the downregulated astrocyte-neuron LR pairs we retrieved the Spon1-App interaction that has previously been shown to block initiating amyloidogenesis (Park *et al*., 2020). Additionally, our analysis also showed an ageing-induced downregulation of the Apoe-Sorl1 interaction which was identified as a downregulated LR pair associated with Alzheimer’s Disease in human prefrontal cortex (Liu *et al*., 2024). As ageing and neurodegenerative disorders are characterized by an early loss of synapses, these altered astrocyte-neuron interactions could underlie ageing-induced synaptic instability (Morrison and Baxter, 2012).

On the other hand, analysis of upregulated astrocyte-microglia interactions similarly revealed ageing-induced changes in multiple GO terms associated with synapse organization and assembly (Fig. 2C). Amongst these altered interactions, we retrieved the Il33-Il1rap LR pair, associated to astrocytic promotion of microglial synapse engulfment and phagocytosis (Vainchtein *et al*., 2018). The dual changes in astrocytes with increased microglia and decreased neuronal signalling make them interesting candidates to further analyse with the aim to discover new and previously missed pathways.

### Transcriptome changes in astrocytes highlight Wnt-signalling

Astrocytes were identified based on their cell type-specific expression of *Sox9*, *Slc1a3*, *Aldh1l1* and *Slc1a2* (Extended Data Fig. 1D). A total of 3383 astrocytes were retrieved, 2138 from adult and 1245 from aged cortices (Fig. 3A). Differential gene expression analysis was performed to identify general molecular differences between adult and aged astrocytes. Over 5000 genes were found to be significantly upregulated in aged astrocytes (p < 0.05 adjusted for multiple comparisons, Extended Data Table 1). Among these, we identified several genes previously reported as upregulated in ageing cortical astrocytes, including *C4b*, *Pcdhb6*, *Pcdhb11*, and *Bmp6* (Boisvert *et al*., 2018). Consistent with the findings of Jin et al., we also observed a significant upregulation of cell–cell adhesion genes, such as *Cdh19*, and ion channel activity genes, including *Grid1* and *Grin2b* (Jin *et al*., 2025). In addition, our dataset also points towards an altered astrocyte synaptic involvement with ageing as expression of synapse elimination genes *Mertk* and *Megf10* were significantly increased (Chung *et al*., 2013; Lee *et al*., 2022).

**Figure 3.**
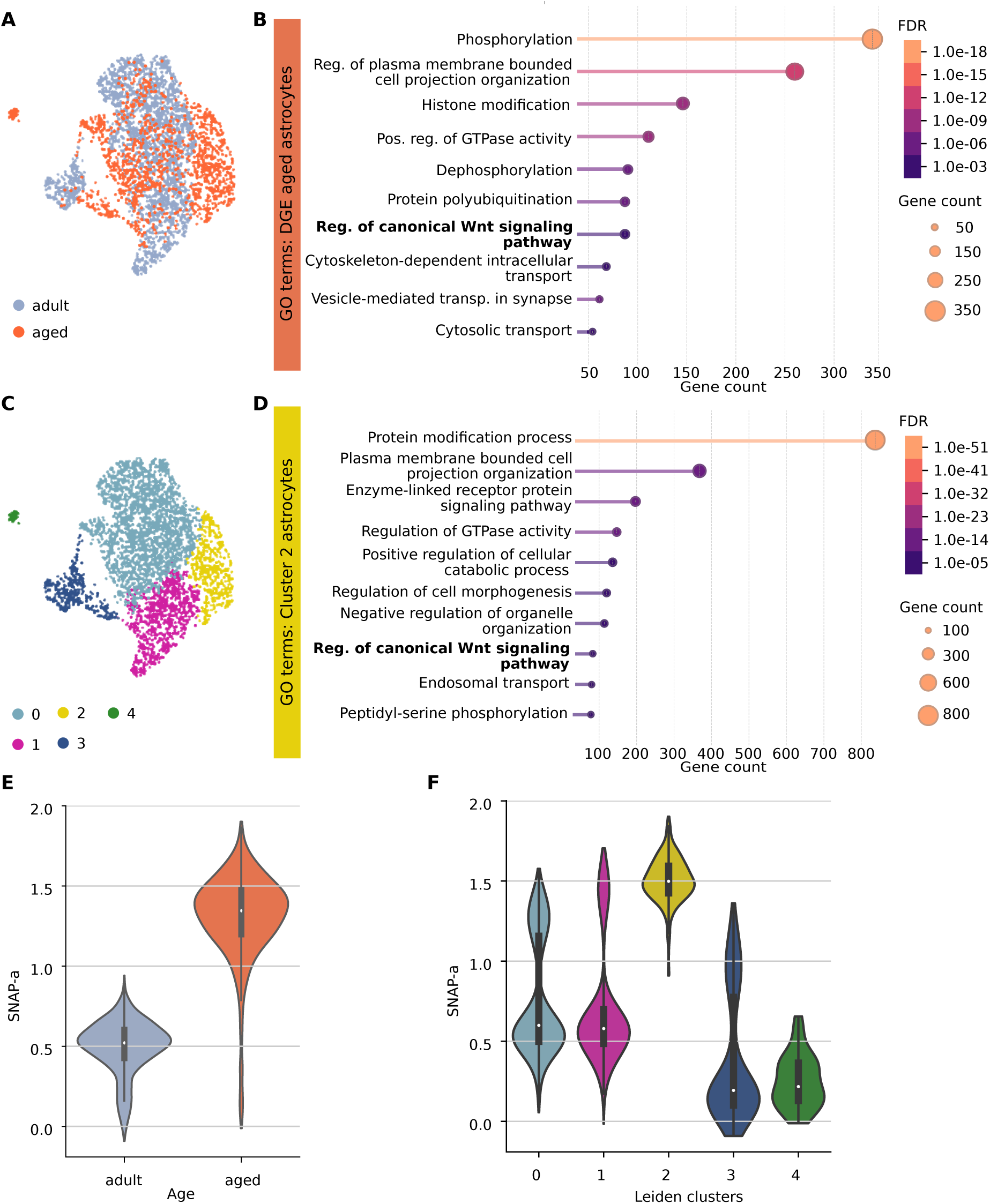
Single cell RNA-seq based profiling of adult and aged cortical astrocytes. **(A)** Umap of astrocyte clusters from adult (colour-coded in grey) and aged (colour-coded in orange) astrocytes. **(B)** STRING-based GO terms of genes significantly enriched in all aged astrocytes. **(C)** Umap of 5 distinct colour-coded astrocyte clusters. **(D)** STRING-based GO terms of genes enriched in astrocyte cluster 2. **(E)** Violin plot of the SNAP-a gene program in aged and adult astrocytes. **(F)** Violin plot of the SNAP-a gene program per astrocytes cluster.

To uncover the molecular pathways affected by ageing in cortical astrocytes, a GO term analysis was performed. Amongst the top GO terms associated with the upregulated genes we found; Histon modification, Protein polyubiquitination, Regulation of canonical Wnt signalling pathway and Vesicle-mediated transport in synapse (Fig. 3B). Amongst the genes associated to the Wnt signalling pathway, we identified genes associated to the regulation of the canonical pathway such as *Tcf7l2*, *Apc* and *Ctnnd2* as well as genes associated to the regulation of the non-canonical pathway including *Daam2*, *Lrp6* and *Wnt5b*. In addition, both positive regulators such as *Lef1* as well as negative regulators such as *Znrf3* and *Wwox* were found. Besides the classical regulators of Wnt signalling, we also identified notch coactivator *Maml2*, necessary for Sox9-dependent Wnt antagonism (Extended Data Table 1) (Sinha *et al*., 2025)

Amongst the significantly downregulated genes (3536) we retrieved genes associated to (Lee *et al*., 2022) proteosome homeostasis such as *Rab5* and *Lamp2*, consistent with the findings of Lee et al that revealed a distinct ageing astrocytic subtype characterized by impaired protein homeostasis (Lee *et al*., 2022)(Extended Data Table 1). Given the regulation of Wnt signalling amongst the up-regulated genes, and as this pathway includes tight feedback regulation, we also looked for targets of the Wnt signalling pathway amongst the significantly down-regulated genes in the aged condition and found the direct Wnt targets *Mmp15*, *Myc* and *Ctnnb1* as well as components of the AP-1 transcription complex, downstream of the non-canonical Wnt signalling pathway, such as *Jun* and *JunD* (Hwang *et al*., 2005; Toualbi *et al*., 2007; J. Liu *et al*., 2022). The top GO terms associated with upregulated genes in adult astrocytes pointed towards Cellular nitrogen compound metabolic process, Mitochondrion organization and Regulation of apoptotic signalling pathway (Extended Data Fig. 2A-B). Together, these results align our dataset with previous reports on ageing astrocytes, including their altered synaptic involvement and impaired protein homeostasis, whilst also revealing a rather unexplored ageing-associated dysregulated pathway namely Wnt signalling.

### Astrocyte cluster-specific changes in Wnt-signalling and their effectors

To explore if dysregulation of the Wnt signalling pathway is a common ageing feature across all cortical astrocytes or would be more pronounced in a specific cluster of astrocytes, a separate re-clustering of all astrocytes was performed (Fig. 3C). This analysis identified five distinct astrocyte clusters: cluster 0 was composed of both adult and aged astrocytes; cluster 1 was predominantly composed of adult astrocytes, and cluster 2 mainly consisted of aged astrocytes. Clusters 3 and 4 were enriched for marker genes of white matter (WM) and layer 1 (L1) astrocytes, respectively (Fig. 3C, Extended Data Fig. 2D-F, (Bocchi *et al*., 2025). Notably, also L1 astrocytes were mostly composed of aged astrocytes, suggesting ageing-specific changes in this layer (Extended Data Fig. 2C, F). However, future experiments will have to collect more of these cells for robust analyses.

Differential gene expression was performed to identify cluster-specific enriched genes (Extended Data Table 2). Among the cluster 2, i.e. enriched in aged astrocytes, we retrieved previously identified Wnt regulators such as *Lrp6, Apc, Daam2, Znrf3, Tcf7l2, Maml2, Ctnnd2, Wwox* and *Lef1* (Extended Data Table 2). Amongst the GO terms associated with cluster 2 enriched genes, we identified protein modification as top GO term, but also again alterations in regulation of canonical Wnt signalling pathway (Fig. 3D). These results suggest that ageing influences Wnt signalling distinctly in cortical astrocyte substates or populations. In addition, expression of the SNAP-a program, previously identified as an astrocyte gene program associated with ageing (Ling *et al*., 2024), was clearly enriched in our aged astrocytes (Fig. 3E) and more specifically in the ageing-associated astrocyte cluster 2 (Fig. 3F). Conversely, among the genes enriched in cluster 1, which is predominantly composed of adult astrocytes, we identified the Jun family transcription factors Jun, JunD, and JunB (Extended Data Table 2). This is line with the hypothesis that Wnt-signalling may be impaired in ageing and hence these downstream effectors of Wnt signalling (Hwang *et al*., 2005; Toualbi *et al*., 2007; Bengoa-Vergniory *et al*., 2014; Arredondo *et al*., 2020) are more highly expressed in the adult astrocytes. GO terms associated with this cluster are among others; Regulation of nucleobase-containing compound metabolic process, Regulation of cell death, Cytoskeleton organization and Neuron projection development (Extended Data Fig. 2C).

### Single-nuclei multiome analysis identifies age-related transcriptional changes including Wnt-signalling and potential up-stream regulators in cortical astrocytes

In order to identify possible up-stream regulators of the changes observed in gene expression above, we performed a multiome analysis of cortical astrocytes. To reliably capture how ageing impacts the local astrocyte population of the mouse cortex GM, an unbiased strategy, without prior astrocyte enrichment, was implemented (Hennes *et al*., 2025). Single nuclei were isolated from the same region and stages as described above (cortical GM of adult (3m-4m) and aged (1y10m-2y) C57BL/6J mice). Quality control analysis selected cells based on gene count (>1000 and <7000) and mitochondrial gene percentage (<5%; Extended Data Fig. 3A-C). This resulted in a dataset comprising 13,778 for RNA-seq and 7965 nuclei for ATAC-seq, respectively (Fig. 4A-B). Cell clustering revealed 7 major cell type clusters, including astrocytes and neurons, that were annotated based on specific cellular markers and external resources (Fig. 4C-D; Extended Data Fig. 3D). Astrocytes were identified based on their cell type-specific expression of *Sox9*, *Slc1a3*, *Aldh1l1* and *Slc1a2* as above (Extended Data Fig. 3E) and subsequently re-clustered. The resulting astrocyte cluster was composed of 808 (ATAC: 366) nuclei, with 273 (ATAC: 110) from adult and 535 (ATAC: 256) nuclei from the aged cortex GM (Fig. 4E-F).

**Figure 4.**
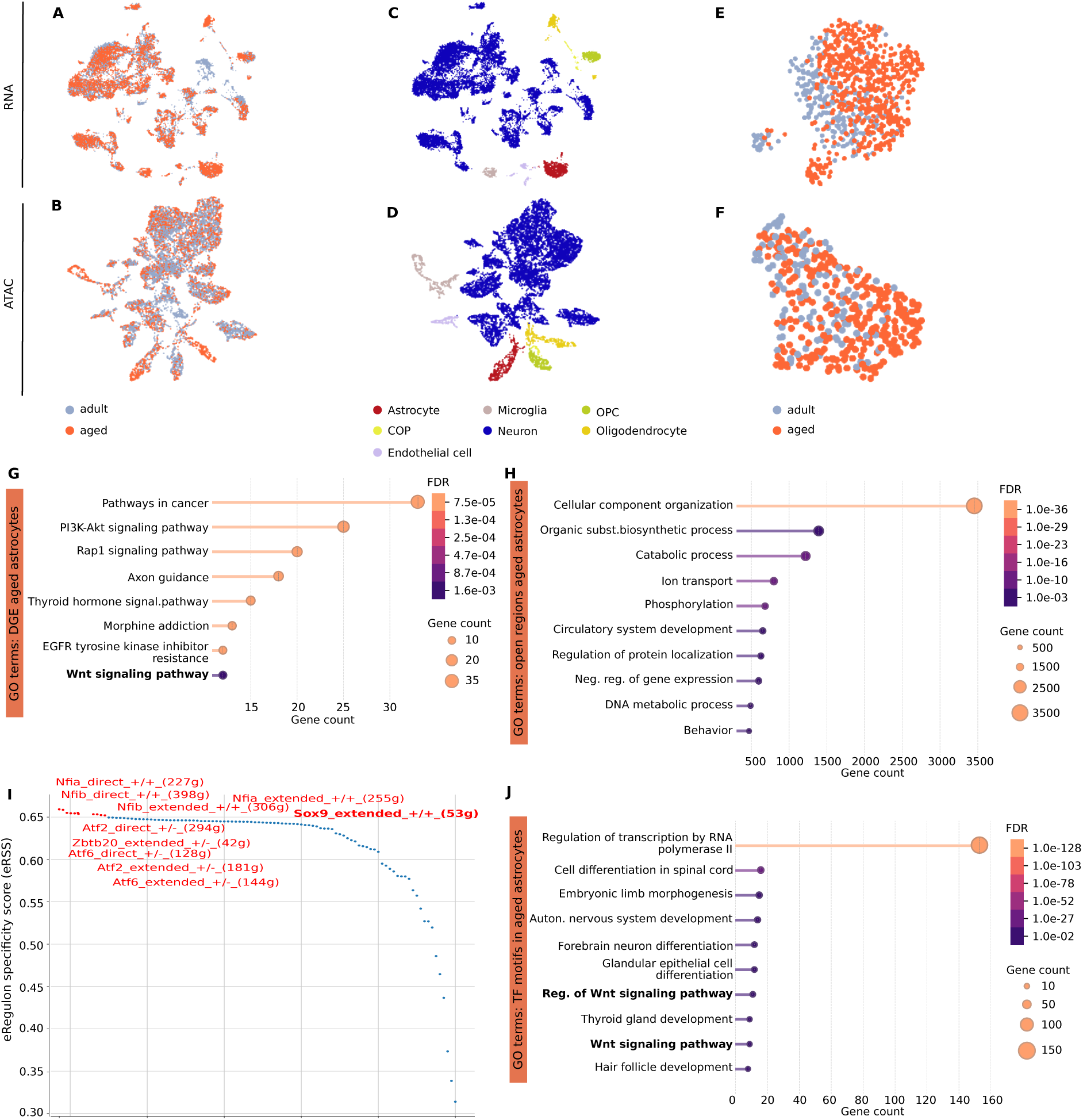
Multiome analysis of adult and aged cortical astrocytes. **(A)-(B)** Umap of RNA (A) and ATAC (B) of 10X Genomics multiome data from adult (colour-coded in grey, n = 1) and aged (colour-coded in orange, n = 1) mouse cortices. **(C)-(D)** Umap of cell cluster annotation of the RNA (C) and ATAC (D) dataset, with cells colour-coded by cell type. COP = committed oligodendrocyte precursor, OPC = oligodendrocyte precursor cell. **(E)-(F)** Umap of astrocyte clusters from adult (colour-coded in grey) and aged (colour-coded in orange) mice for the RNA (E) and ATAC dataset (F). **(G)** STRING-based GO terms of genes significantly enriched in aged vs. adult astrocytes. **(H)**. STRING-based GO terms of genes associated to open regions in aged astrocytes. **(I)** SCENIC+ specificity score of eRegulons consisting of enriched motifs and their associated transcription factors that are more specific to aged versus adult astrocytes. **(J).** STRING-based GO terms of transcription factor motifs in ageing astrocytes.

Differential gene expression analysis was performed on the snRNA-seq dataset and differential peak calling was implemented on the ATAC-seq dataset to identify differences in accessible chromatin regions in aged astrocytes (Extended Data Tables 3 and 4). GO term analysis of significantly upregulated genes indicated the top pathways altered in ageing astrocytes such as; Pathways in cancer, PI3K-Akt signalling pathway, Rap1 signalling, Axon guidance, Thyroid hormone signalling pathway, Morphine addiction, EGFR tyrosine kinase inhibitor and Wnt signalling (Fig. 4G), in line with the scRNA-seq shown above. On the other hand, GO term analysis of the genes associated to the differentially open regions in aged astrocytes pointed towards more general pathways such as; Catabolic process, Ion transport, Phosphorylation, Regulation of protein localization and Negative regulation of gene expression (Fig. 4H). Notably, among the significant GO terms we again identified Regulation of Wnt signalling (Extended Data Fig. 3F).

A joint profiling of chromatin accessibility and gene expression analysis was achieved using SCENIC+ to reveal enhancer-driven gene regulatory networks (González-Blas *et al*., 2023). SCENIC+ identified eRegulons consisting of enriched motifs and their associated transcription factors (TFs) in adult and aged astrocytes, revealing active TF regulatory networks, including those driven by *Sox9*, *Nfib*, and *Nfia* (Fig. 4I) to be more specific for aged astrocytes. GO analysis of enriched motifs in ageing astrocytes identified pathways such as; Regulation of transcription by RNA polymerase 2, Cell differentiation in spinal cord, Autonomic nervous system development, Forebrain neuron differentiation and Regulation of the Wnt signalling pathway (Fig. 4J, Extended Data Table 5). Thus, both our datasets converge on a novel pathway, so far not detected in ageing astrocytes, Wnt signalling and its down-stream transcriptional effectors (including AP1 TFs).

### A conserved ageing-associated astrocyte cluster shows dysregulation of Wnt signalling across species

To ensure the relevance of Wnt signalling across species, we performed additional astrocyte sub-clustering analysis, using the same parameters as for our scRNA-seq dataset (Fig. 5A), on our multiome dataset (Fig. 5B) as well as a human snRNA-seq dataset (Fig. 5C). The latter was composed of snRNA-seq data from healthy young (23-30 years old) and aged (70-74 years old) human prefrontal cortices (Chien *et al*., 2024). The dataset included 39,830 nuclei with an average of 213,000 RNA reads and 6800 genes detected per cell. Heatmap visualization of the percentage overlap of differentially expressed genes between these different astrocyte clusters revealed that our ageing-associated astrocyte cluster 2 is indeed transcriptionally similar to the ageing-enriched astrocyte cluster 1 in the multiome and cluster 0 in the human dataset (Fig. 5D). GO terms associated with the shared genes between cluster 2 and human cluster 0 are; Synapse organization, Regulation of autophagy and Regulation of Wnt signalling pathway (Fig. 5E). Among the shared genes between mouse and human aged astrocytes, we retrieved also the previously identified Wnt candidates; *Tcf7l2*, *Maml2* and *Wwox* (Extended Data Table 6). Intriguingly, also the AP1 transcription factors of the JUN family were higher in the human astrocytes from the younger brains (Extended Data Table 6). These results thus indicated that physiological ageing induces a dysregulation of the Wnt signalling and Jun transcription factors in cortical astrocytes across species.

**Figure 5.**
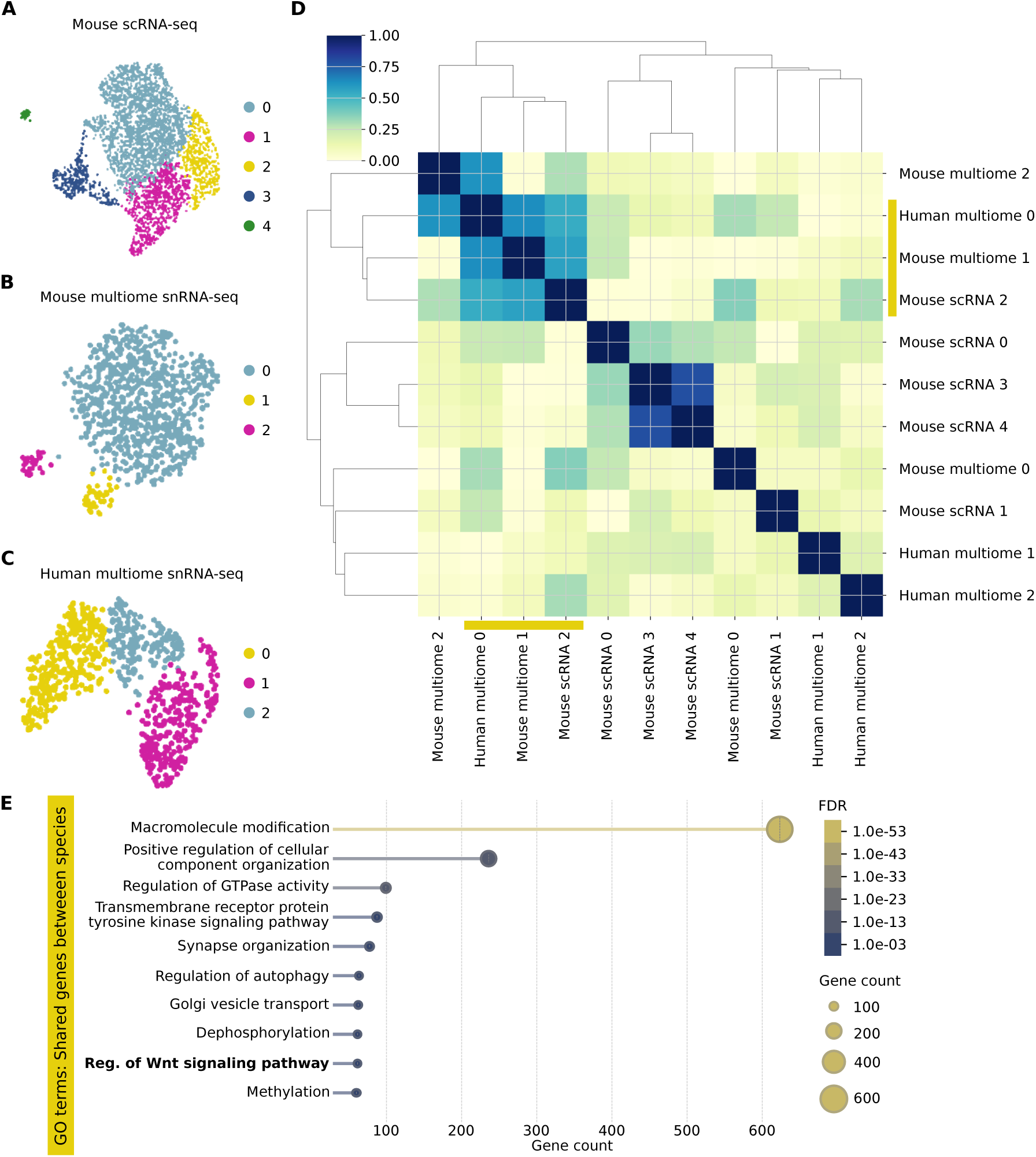
A conserved ageing astrocyte cluster shows Wnt dysregulation. **(A)** Umap of scRNA-seq astrocyte clusters. **(B)** Umap of snRNA-seq astrocyte clusters obtained from the multiome dataset. **(C)** Umap of astrocyte clusters from a human snRNA-seq dataset of young and old patients. **(D).** Heatmap of the percentage overlap of differentially expressed genes between different astrocyte clusters across datasets (t-test, Benjamini-Hochberg adjusted p-value < 0.05 in all datasets). **(E)** STRING-based GO terms of genes shared between single cell astrocyte cluster 2 from (A) and human astrocyte cluster 0 from (C).

### Ageing alters the expression levels of Wnt signalling components

To verify the ageing effect on astrocytic Wnt signalling, Wnt candidates were selected for further investigation. Dot plot visualization of gene expression levels of the identified Wnt candidates upregulated (Fig. 6A) or downregulated (Fig. 6B) in ageing revealed clear differences for *Daam2*, *Tcf7l2*, *Znrf3*, *Maml2*, *Wwox*, *Ctnnd2*, *Apc* and *Ctnnb1* and *Lef1*, *Lrp6* and *Wnt5b* (Fig. 6A-B). Feature plots were created to explore if the candidate upregulation is cluster-specific or common to all aged astrocytes (Fig. 6C).

**Figure 6.**
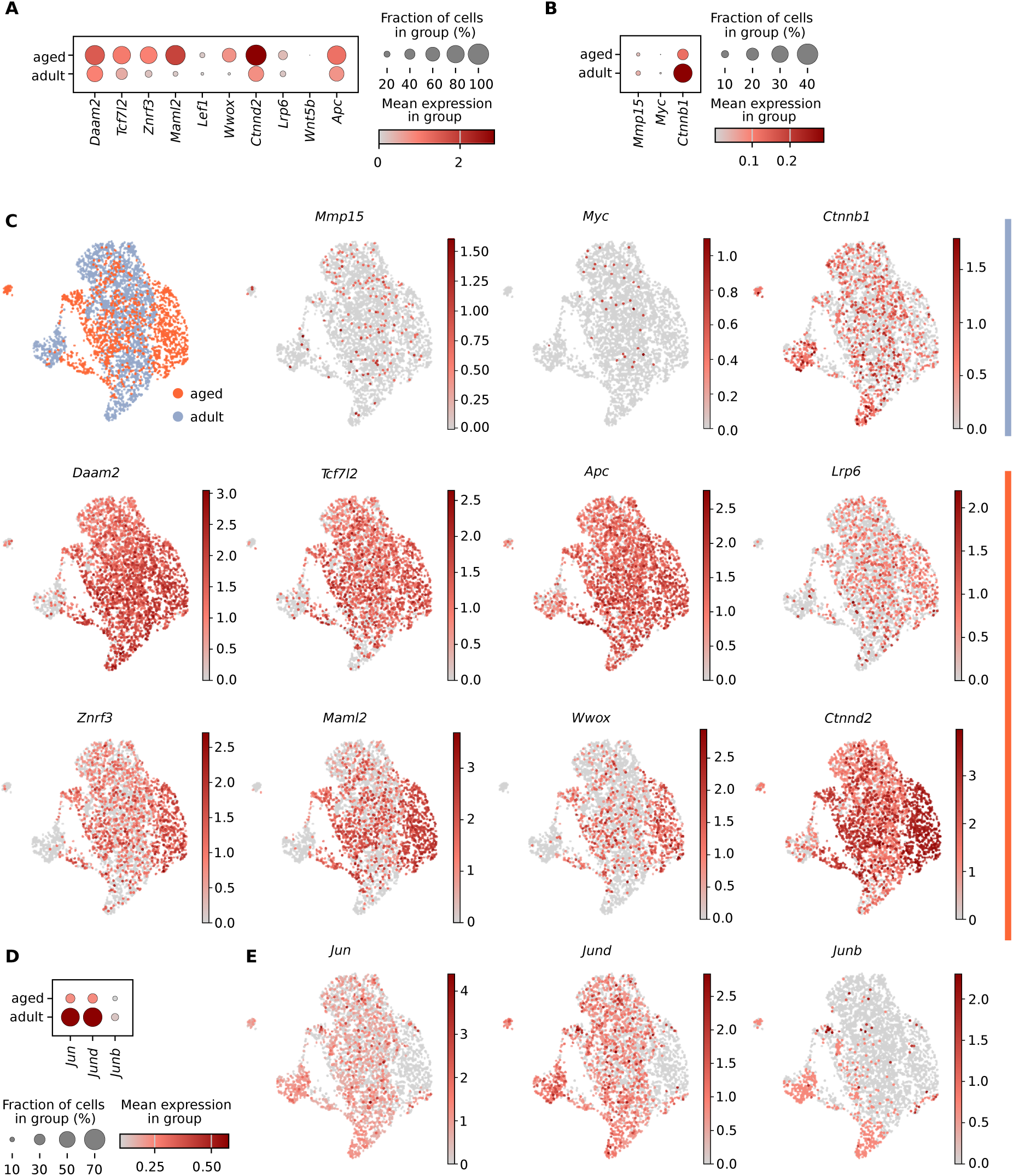
Altered gene expression levels of Wnt signaling candidates in ageing astrocytes. **(A)** Dot plot illustrating the expression of ageing-upregulated Wnt pathway genes (t-test, Benjamini-Hochberg adjusted p-value < 0.05). **(B)** Dot plot revealing expression levels of ageing-downregulated Wnt pathway genes (t-test, Benjamini-Hochberg adjusted p-value < 0.05). **(C)** Umap of Wnt candidate genes across astrocyte clusters. **(D)** Dot plot of Jun family transcription factors. **(E)** Umap illustrating Jun family transcription factor expression across astrocyte clusters.

Among the selected Wnt regulators identified as upregulated in aged astrocytes; *Wwox*, *Znrf3*, *Ctnnd2* and *Maml2* showed the most pronounced differential expression patterns and were seemingly highest expressed in ageing astrocyte cluster 2 (Fig. 6C).

Amongst the Wnt pathway candidates identified as downregulated in aged astrocytes, *Myc* and *Mmp15* seemed to be very heterogeneous in adult astrocytes, while *Ctnnb1* was more widespread in adult astrocytes and hence exhibited a more distinct differential expression compared to the aged astrocytes (Fig. 6C). Taken together, these differential expression patterns of Wnt regulators show mostly upregulation of Wnt inhibitors and downregulation of Wnt effectors, suggesting that Wnt signalling seems decreased in aged astrocytes.

To further probe this concept, we examined AP-1 transcription-complex components (Jun, JunD and JunB), which are amongst the downstream effectors of Wnt signalling (Hwang *et al*., 2005; Toualbi *et al*., 2007; Bengoa-Vergniory *et al*., 2014; Arredondo *et al*., 2020) and were downregulated in aged astrocytes (Fig. 6D). Dotplot representation, as well as feature plots clearly visualized the significant downregulation of *Jun* and *JunD* in aged astrocytes (Fig. 6D-E). Moreover, the cluster 2 aged astrocytes associated with dysregulated Wnt signalling showed an almost complete loss of *Jun*, *JunD*, and *JunB* expression. These data suggest a notable down-regulation of the Wnt-signalling pathway and some of its effectors, which promoted us to explore this at the protein and functional level.

### RNAscope and immunohistochemical profiling of Wnt pathway and effectors in the ageing cortex

To further validate the above findings and explore if layer-specificity contributes to the observed heterogeneity, we explored *Maml2* and *Daam2* RNA levels by RNAscope in coronal sections of C57BL/6J adult (3-4m) and aged (22-24m) cerebral cortices. RNAscope probes for the Wnt genes were co-applied with a *Slc1a3* probe for the astrocyte-specific glutamate transporter Glast to restrict the analysis specifically to astrocytes. Figure 7A shows representative images of cortical bins of adult and aged brain sections used for quantification of *Maml2*. Using QuPath, we analysed the percentage of *Slc1a3^+^*astrocytes also positive for the *Maml2* signal (≥ 3 dots/cell) as well the percentage of *Maml2^+^/Slc1a3^+^* astrocytes with high *Maml2* signal (≥ 10 dots/cell). Notably, about half of all astrocytes were *Maml2^+^* and this proportion is not statistically different between adult and aged conditions (Fig. 7B). However, in the aged condition, a significantly larger proportion of *Slc1a3*^+^ astrocytes exhibit a high (≥ 10 dots/cell) *Maml2* signal (Fig. 7C).

**Figure 7.**
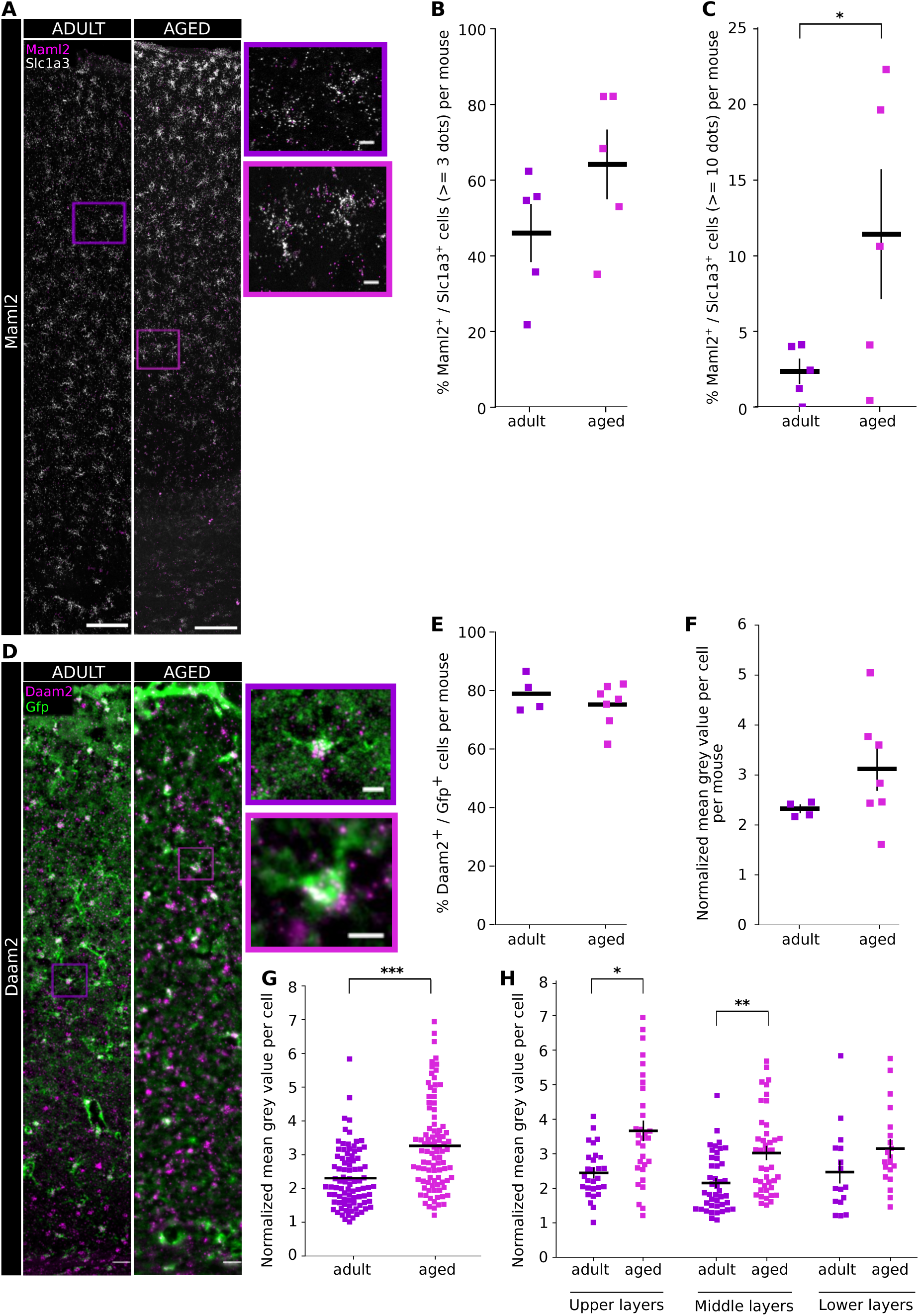
RNAscope analysis of Wnt candidates in aged cortical astrocytes. **(A)** Representative images of cortical bins of adult and aged coronal brain sections used for *Slc1a3* and *Maml2* RNA-scope analysis. Magenta = *Maml2*, white = *Slc1a3*, scale bar = 100µm, inserts= 10µm purple (adult), pink (aged). **(B)** Analysis of *Slc1a3⁺* astrocytes positive for *Maml2* (≥3 RNA dots per nucleus) showed no significant difference between adult and aged mice. n = 5 per condition **(C)** Percentage of *Slc1a3⁺* astrocytes with higher *Maml2* expression (≥10 RNA dots per nucleus) was significantly increased in aged astrocytes. Unpaired t-test (one-sided), p-value = 0.0325. **(D)** Representative images of cortical bins of adult and aged Aldh1l1-eGFP coronal brain sections used for *Daam2* RNAscope analysis. Magenta = *Daam2*, green = eGFP, scale bar= 20µm, inserts= 10µm purple (adult), pink (aged). **(E)** Percentage of *Daam2^+^*/GFP^+^ cells is not different between adult (n = 4) and aged (n = 7) astrocytes. **(F)** No difference can be observed analyzing normalized mean grey values of *Daam2^+^* single dots/ nuclei between adult and aged astrocytes grouped per mouse. **(G)** *Daam2* signal intensity, measured as normalized mean grey value, is significantly increased in aged cortical astrocytes. Mann Whitney test, p-value= 0.0004 **(H)** Layer distribution analysis reveals that the increase in Daam2 signal intensity is attributed to astrocytes in the upper and middle but not lower layers. Upper layers: unpaired t-test, p-value= 0.03; middle layers: Mann Whitney test, p-value=0.0033.

For *Daam2*, we used an alternative method to confirm astrocytic co-localization and adapted our analysis strategy, as the high density of *Daam2* puncta per cell precluded reliable segmentation and quantification of individual signals by QuPath. *Daam2* RNAscope probes were thus applied to coronal sections of Aldh1l1-eGFP adult (3-4m) and aged (22m-24m) mice followed by immunostaining for GFP. Figure 7D shows a representative image of bins spanning the entire thickness of adult and aged cortical GM used for quantification of *Daam2* RNAscope signal. A high proportion of Aldh1l1-eGFP^+^ astrocytes had *Daam2* signal (about 80%) with no difference in this proportion detectable between adult and aged conditions (Fig. 7E). When the normalized mean grey value of *Daam2*⁺/*Aldh1l1*⁺ cells was analysed at the per-animal level, we observed a consistent trend toward higher signal intensity in aged mice that did not reach statistical significance due to substantial inter-animal variability (Fig 7F). In contrast, single-cell–level analysis revealed a significantly higher signal in the aged condition (Fig. 7G).

When examining the layer distribution, we noted that mostly astrocytes in upper and middle but not lower layers exhibited a higher *Daam2* expression in aged cortices (Fig. 7H).

### Immunohistochemical analysis of AP1 transcriptome components in aged astrocytes

Next, we stained for Jun, JunD or JunB that were down-regulated at mRNA level in aged astrocytes (Fig. 8A) together with the astrocyte marker S100□ (see e.g. (Ohlig *et al*., 2021; Bocchi *et al*., 2025)) in coronal sections of adult (3-4m) and aged (22m-24m) cortical GM. Figure 8A shows representative images indicating the bins used for quantification and higher magnification images to visualize how double positive cells were selected. The percentage of Jun^+^/S100b^+^ astrocytes showed only a trend towards downregulation in the aged condition (Fig. 8B), while the percentage of both JunD^+^ and JunB^+^ astrocytes were significantly decreased in the aged cortex (Fig. 8C-D).

**Figure 8.**
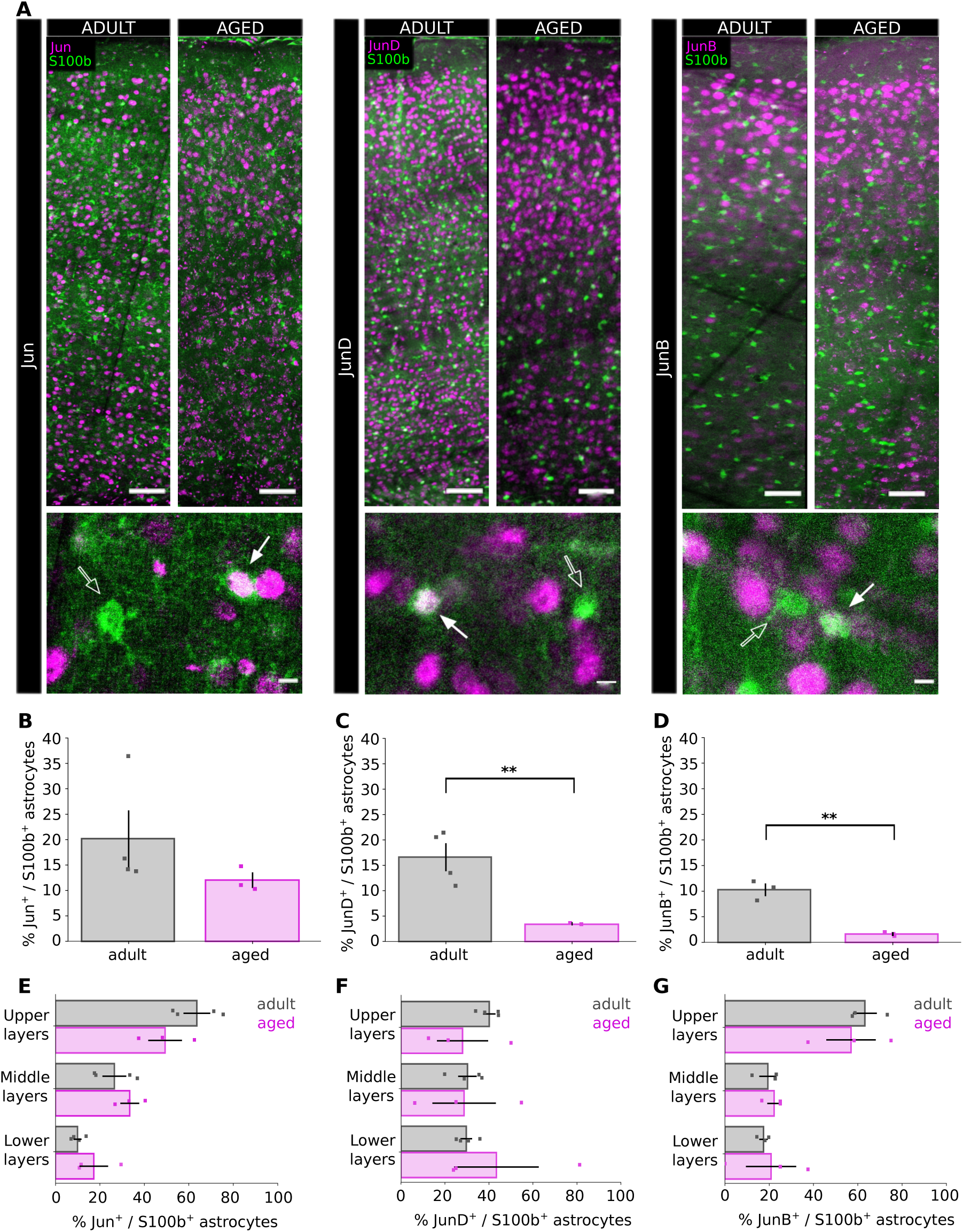
Immunohistochemical characterization of Jun transcription factors in adult and aged cortical astrocytes. **(A)** Representative images of cortical bins of adult and aged coronal brain sections used for analysis of Jun, JunD and JunB. Magenta = Jun/JunD/JunB, green= S100b, scale bar: 50µm and 5µm (inserts), white arrow = positive signal, empty arrow = negative signal. **(B)** Statistical analysis of the percentage of Jun^+^/S100b^+^ astrocytes in adult (n = 4) and aged (n = 3) mice revealed no difference. **(C)** Statistical analysis of the percentage of JunD^+^/S100b^+^ astrocytes in adult (n = 4) and aged (n = 3) mice showed a significant decrease of JunD^+^/S100b^+^ in aged astrocytes. Unpaired t-test, p-value = 0.0076. **(D)** Statistical analysis of JunB^+^/S100b^+^ astrocytes in adult (n = 3) and aged (n = 3) mice revealed a significant decrease in JunB^+^/S100b^+^ aged astrocytes. Unpaired t-test, p-value= 0.0015. **(E)-(G)** Layer distribution analysis showed no differences in the percentage of Jun^+/^S100b^+^ (E), JunD^+^/S100b^+^ (F) and JunB^+^/S100b^+^ (G) astrocytes.

As we noted that Jun transcription factors are present only in subsets of astrocytes, we aimed to explore if this was related to specific layers, by taking advantage of a dataset, in which we had used the Fezf2-transgenic layer marker to dissect layers 2-4, 5 and 6 (see Methods and (Stogsdill *et al*., 2022) followed by magnetic-associated cell sorting (MACS) of astrocytes as described (Ohlig *et al*., 2021). As no myelin removal could be done given the cell number limitations, oligodendrocytes were included besides astrocytes in the dataset (Extended Data Fig. 4) as previously described (Ohlig *et al*., 2021). When focussing on astrocytes, 7 clusters were detected (Extended Data Fig. 5A, B) with the furthest distant cluster, cluster 5, probably reflecting astrocytes in or close to the White Matter (WM), given the high score based on our previous WM astrocyte isolation (Extended Data Fig. 5C; (Bocchi *et al*., 2025)). Most of the clusters contained astrocytes from several layers, but cluster 1 consisted almost exclusively of Layer2-4 astrocytes (Extended Data Fig. 5B). The heatmap in Extended Data Fig. 5D shows the top 12 differentially expressed genes between astrocytes from the different layers, such as e.g. Chrodin-like higher in Layer2-4 astrocytes consistent with previous data (Lanjakornsiripan *et al*., 2018). Interestingly, we noted several members of the immediate early gene (IEG) transcription factor family (such as *Fos*, *JunB*, *Jun*, *Ier2, Egr1*) expressed at higher levels in astrocytes of Layer2-4 (Extended Data Fig. 5D-F, Extended Data Table 7). This prompted us to examine the layer distribution of Jun+ astrocytes at the protein level and explore if this would change in ageing. Layer distribution analysis indeed revealed a higher proportion of Jun^+^ and JunB^+^, but not JunD^+^, astrocytes in the upper layers (Fig. 8E-G), thereby also confirming the transcriptome data. Notably, however, this distribution was not changed in the aged cortex, suggesting that astrocytes in all layers down-regulate these transcription factors retaining more double^+^ astrocytes in the upper layers (Fig. 8E-G).

### JunD overexpression analysis in aged astrocytes

As the above transcriptome and immunohistochemical analysis revealed a significant downregulation of JunB/D expression in aged astrocytes, we aimed to explore the consequences of reinstalling their expression in aged astrocytes. To target astrocytes specifically we used a Mokola-pseudotyped lentivirus previously shown to target astrocytes in the adult cerebral cortex (Natarajan *et al*., 2024). Viral vectors expressing mScarlet alone (control) or co-expressing JunD and mScarlet were injected into the cortical GM of aged mice and analysed one week later. As the viral vector injection generates a mild injury (Natarajan *et al*., 2024), and a sign of ageing of reactive astrocytes is their reduced proliferation (Heimann *et al*., 2017), we probed proliferation of transduced aged astrocytes by providing the thymidine analogue 5-ethynyl-2’-deoxyurdine (5-EdU) in drinking water for one week to the mice. Figure 9A shows a representative image of cortical bins, from reporter control- and a JunD-vector injected cortical GM sections, immunostained for JunD/EdU and RFP. Analysis of Sox9^+^/RFP^+^ cells confirmed astrocyte specificity (Fig. 9B). Mice injected with the JunD-vector showed a clear visible increased JunD signal in the RFP^+^ transduced astrocytes that was absent or lower in the reporter control-vector condition (Fig. 9A). We next analysed the amount of EdU^+^/RFP^+^ astrocytes in both conditions and found a significant increase in proliferating EdU^+^ astrocytes in the JunD overexpression condition (Fig. 9C). We also stained for Hmgb1, as loss of its nuclear expression has been linked to senescence and ageing (Gaikwad *et al*., 2021; Sofiadis *et al*., 2021; Salminen, 2026) and its systemic application has been used to counteract ageing and pathology (Komkleow *et al*., 2026; Supasai *et al*., 2026). Moreover, the family member Hmgb2 labels proliferating astrocytes in the WM (Bocchi *et al*., 2025) and both family members have also been found downregulated in our aged single cell dataset. Indeed, JunD re-expression significantly increased the percentage of Hmgb1^+^/RFP positive cells compared to controls (Fig. 9D). Taken together, these results suggest that increasing the levels of JunD in aged astrocytes reinstalls some aspects of a more youthful state with higher Hmgb1 levels counteracting ageing, senescence, and oxidative challenges accompanied by increased proliferation in aged astrocytes.

**Figure 9.**
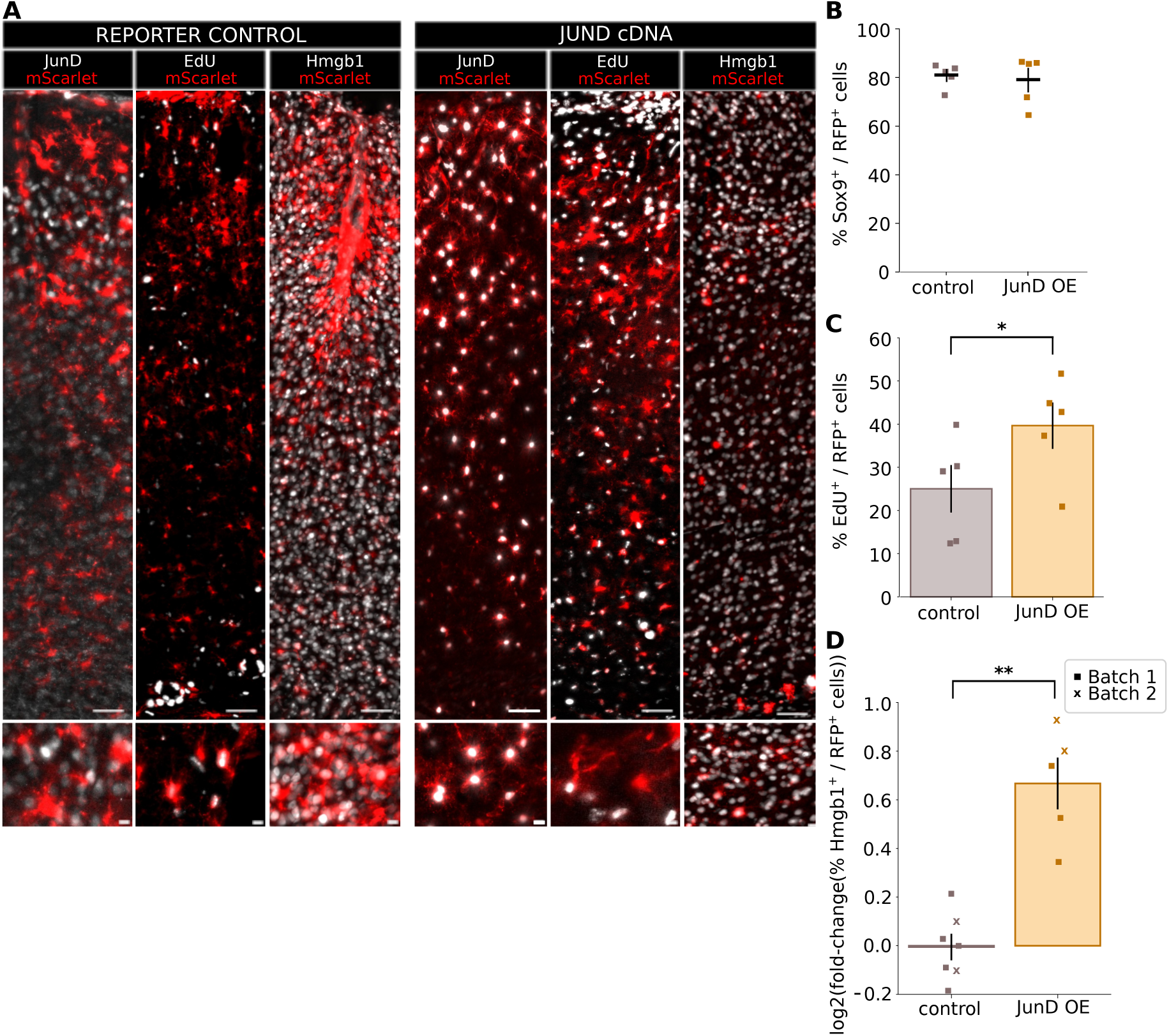
JunD cDNA overexpression in aged astrocytes reinstates a more youthful proliferative state. **(A)** Representative images of cortical bins of aged mice injected with either reporter control vector (left) or JunD cDNA vector (right) immunostained for JunD, EdU, Hmbg1 and RFP. Red = RFP/mScarlet, white = JunD, EdU, Hmgb1, scale bar = 50µm and 10µm (inserts). **(B)** Analysis of Sox9^+^/RFP^+^ cells indicates that around 80% of the targeted cells are astrocytes in both reporter control (n = 5) and JunD cDNA (n = 5) vector injected mice. **(C)** Statistical analysis showed a significant increase in the percentage of EdU^+^/RFP^+^ cells in mice injected with the JunD cDNA vector. Unpaired t-test, p-value = 0.043. n = 5 per condition **(D)** Log2-fold change of the percentage of Hmgb1^+^/RFP^+^ cells against mean control significantly increases in mice injected with the JunD cDNA vector. Unpaired t-test with unequal variance, p-value = 0.001148. n = 7 mice (control), 5 mice (JunD OE).

## Discussion

Ageing is a complex, multifactorial biological process that differentially impacts distinct cellular populations, ultimately leading to functional decline. In the central nervous system, deciphering the effects of ageing has been particularly challenging due to the heterogeneity of the different cell types (Siletti *et al*., 2025). Indeed, our transcriptome analysis demonstrates that ageing elicits both quantitatively and qualitatively distinct responses in different cell types and their subtypes, not only between neurons and glial cells but also across different glial cell populations and amongst astrocytes e.g. mostly in one cluster. Further investigation of how ageing-related changes affect intercellular communication revealed increased astrocyte–microglia signalling and reduced astrocyte–neuron interactions, which could both impact on synaptic homeostasis that is disturbed in ageing. These dual alterations in astrocytes highlighted them as compelling candidates for a key role in brain ageing.

So far, studies exploring astrocytes in ageing already indicated a switch in the astrocyte population towards a more inflammatory phenotype, suggesting they primarily contribute to ‘inflammageing’ that progresses cognitive deterioration (Ximerakis *et al*., 2019; Allen *et al*., 2023; Hahn *et al*., 2023). Studies focused particularly on ageing astrocytes, in addition revealed functional and morphological alterations that impact their ability to maintain synaptic homeostasis (Boisvert *et al*., 2018; Popov *et al*., 2021; Lee *et al*., 2022).

By implementing an integrated multi-omics approach, we revealed a previously unnoticed pathway deregulated in ageing astrocytes, namely Wnt signalling. The Wnt signalling pathway is well known for its fundamental roles during development in regulating cell fate, proliferation, and tissue patterning (Xue *et al*., 2025). How the activity of the Wnt signalling pathway shifts upon ageing—and how these age-related alterations differentially affect distinct cell types—remains much less understood. Interestingly, downregulation of Wnt signalling dynamics have been described in other ageing animal models such as the Ocotodon degus and the African turquoise killifish (Inestrosa *et al*., 2020; Ogamino *et al*., 2024). In mice, previous work examining inter-regional communication showed that the isocortex influences other brain regions through Wnt-mediated signalling, and this regulatory interaction becomes disrupted with ageing (Wu *et al*., 2024). Moreover, Alzheimer’s disease downregulates neuronal Wnt signalling, leading to synaptic vulnerability (Flores *et al*., 2024). Notably, ageing OPCs display upregulated Wnt signalling that impairs their differentiation capacity (Heo *et al*., 2025), highlighting that age-associated changes in Wnt pathway activity are cell-type-specific and are not uniformly manifested across the brain.

Our analysis of aged cortex GM astrocytes indicated an upregulation of regulators and effectors of Wnt signalling, the majority of which exert an overall inhibitory influence on Wnt pathway activity. SCENIC+ analysis revealed that TF drivers in aged astrocytes are involved in the regulation of the Wnt pathway. Among these TF drivers, we identified *Sox9* that has previously been shown to antagonize Wnt signalling by inducing expression of Maml2, capable of affecting β-catenin turnover (Sinha *et al*., 2025). *Maml2* was upregulated in aged astrocytes in our dataset and validated via RNAscope in aged mouse cortex. In addition, we also identified and validated an increase in *Daam2* expression in aged astrocytes. Daam2 (Dishevelled associated activator of morphogenesis 2) belongs to the family of Formin proteins and acts as a context-dependent modulator of Wnt signalling by its interaction with Dishevelled. Although Daam family proteins classically promote Wnt signalling, Daam2 suppresses canonical Wnt/β-catenin activity in oligodendrocyte lineage cells by Dishevelled-dependent mechanisms (Wang *et al*., 2023). Interestingly, in astrocytes, loss of Daam2 led to a reduction in astrocytic morphological complexity thereby impairing synaptic function (Jo *et al*., 2021).

In line with the evidence for reduced Wnt signalling, we observed and validated the down-regulation of members of the AP1 transcriptome complex such as Jun and JunD, also functioning downstream of Wnt signalling (Hwang *et al*., 2005). C-Jun is a direct target of the b-catenin/Tcf4 transcription complex regulated by Wnt (Saadeddin *et al*., 2009) and is also regulated by Wnt3a (Hwang *et al*., 2005; Gujral and MacBeath, 2010). Collectively, our data indicate that ageing is associated with reduced Wnt signalling activity in astrocytes. Given that astrocyte communication with microglia and endothelial cells—shaping processes such as synapse elimination and blood–brain barrier maintenance—is mediated in part by Wnt signalling, age-related downregulation of this pathway could contribute to multiple ageing-associated phenotypes in the brain (Guérit *et al*., 2021; Faust *et al*., 2025). Ageing-related deregulation of Wnt signalling in astrocytes may thus impair their communication with other cell types, thereby increasing synaptic vulnerability, a process that fundamentally underlies both physiological and pathological cognitive decline. Importantly, these findings are conserved across species, as a human ageing dataset revealed both an ageing-associated astrocyte specific cluster with comparable dysregulation of Wnt signalling and higher expression of JUN transcription factor family members in younger astrocytes.

A remarkable, yet often underappreciated, feature of astrocytes is their plasticity—the capacity to continuously adapt to a changing environment. Interestingly, it was recently shown that astrocytes can switch between states, that the same astrocyte can be neuroprotective before becoming neurotoxic (Zhang *et al*., 2025). It is therefore interesting to hypothesize that also the astrocyte ‘states’ in the ageing brain are not fixed, but can be influenced and potentially be reversed to more ‘youthful’ states. We demonstrated this by re-expressing JunD as we had identified a significant downregulation of JunD in aged astrocytes, most pronounced in our ageing-associated astrocyte cluster that also showed most deregulated Wnt signalling. JunD, as an AP-1 complex member, can activate or repress transcriptional responses involved in cell proliferation, differentiation and death (Shaulian and Karin, 2002). Interestingly, downregulation of JunD has previously been reported in liver ischemia–reperfusion injury, endothelial cells, and cochlear hair cells, where it has been associated with a loss of protection against oxidative stress, senescence, and apoptosis (Weitzman *et al*., 2000; Paneni *et al*., 2013; H. Liu *et al*., 2022). Thus, the newly discovered reduction of Jun transcription factors in ageing astrocytes may relate to these functions. Importantly, re-expressing JunD in aged cortical astrocytes was capable of improving their proliferation upon a mild injury, indicated by increased EdU-incorporating astrocytes. This is promising as astrocyte proliferation in response to injury is greatly reduced in the ageing brain (Heimann *et al*., 2017). Moreover, we observed effects on Hmgb1-expression, the reduced levels of which are a sign of senescence, and whose multiple intra- and extracellular functions implicated it in many aspects of inflammageing (Komkleow *et al*., 2026; Salminen, 2026; Supasai *et al*., 2026). This shows that at least as monitored by two phenotypes, aged astrocytes can be triggered to reacquire a more juvenile phenotype that could potentially promote healthy brain ageing. However, a more comprehensive analysis of the phenotype of the JunD-re-expressing astrocytes is required to fully understand their “rejuvenation”.

## Conclusion

By using a multi-omics approach to characterize ageing astrocytes at multiple molecular levels, we identified a novel conserved ageing-affected pathway in cortical astrocytes, namely Wnt signalling and the downstream Jun transcription factors. Our results suggest that this dysfunction could drive other known ageing-induced astrocyte impairments. By reintroducing JunD in aged astrocytes, we show that some aspects of the aged astrocyte states can be reversed, which further suggests to exploit the astrocyte plasticity for therapeutic approaches to combat physiological and pathological brain ageing.

## Acknowledgements

We would like to thank Tatiana Simon (BMC) for her help with the sc/snRNA-seq library preparation, Ines Muehlhahn for her support with cloning (BMC) and Paulina Chlebik (BMC, Munich) for her help with lentiviral production and titration. We thank the Core Facility Genomics at Helmholtz Munich for their excellent consultation and sequencing, the Core Facility Bioimaging (BMC) for their excellent support. We acknowledge the support and assistance of Tobias Straub from the Bioinformatics Core facility (BMC). We sincerely thank all members of the Götz lab for their valuable insights and contributions throughout this project. This study was supported by funding from the Helmholtz Association and the following funding to M.G.: The Initiative and Networking Fund of the Helmholtz Association, the European Union (NSC Reconstruct project 874758 and advanced ERC NeuroCentro project 885382), the German Research Foundation (SyNergy project 390857198, TRR274 project 408885537, Immunostroke project 40535880) and a New Frontiers in Research Fund Transformation grant, funded through three Canadian federal funding agencies (CIHR, NSERC, and SSHRC).

## Author Contributions

M.G. and M.H. conceived and designed the project. M.H., M.L.R. and J.F.-S. contributed to shaping the project. M.H. performed all experiments with assistance of M.T. for the animal experiments and RNAscope, except the isolation of astrocytes from different layers performed by J.A.S. and M.G. under supervision of P.A. J.F.-S. provided the adult cortical grey matter scRNA-seq dataset. C.L.L. designed, produced and validated the lentiviral vectors used in this study. M.L.R. performed all bioinformatic analysis and support. M.H., M.L.R., J.F.-S. and M.G. wrote the manuscript. M.G. provided all the funding.

---

**Extended Data Figure 1.**
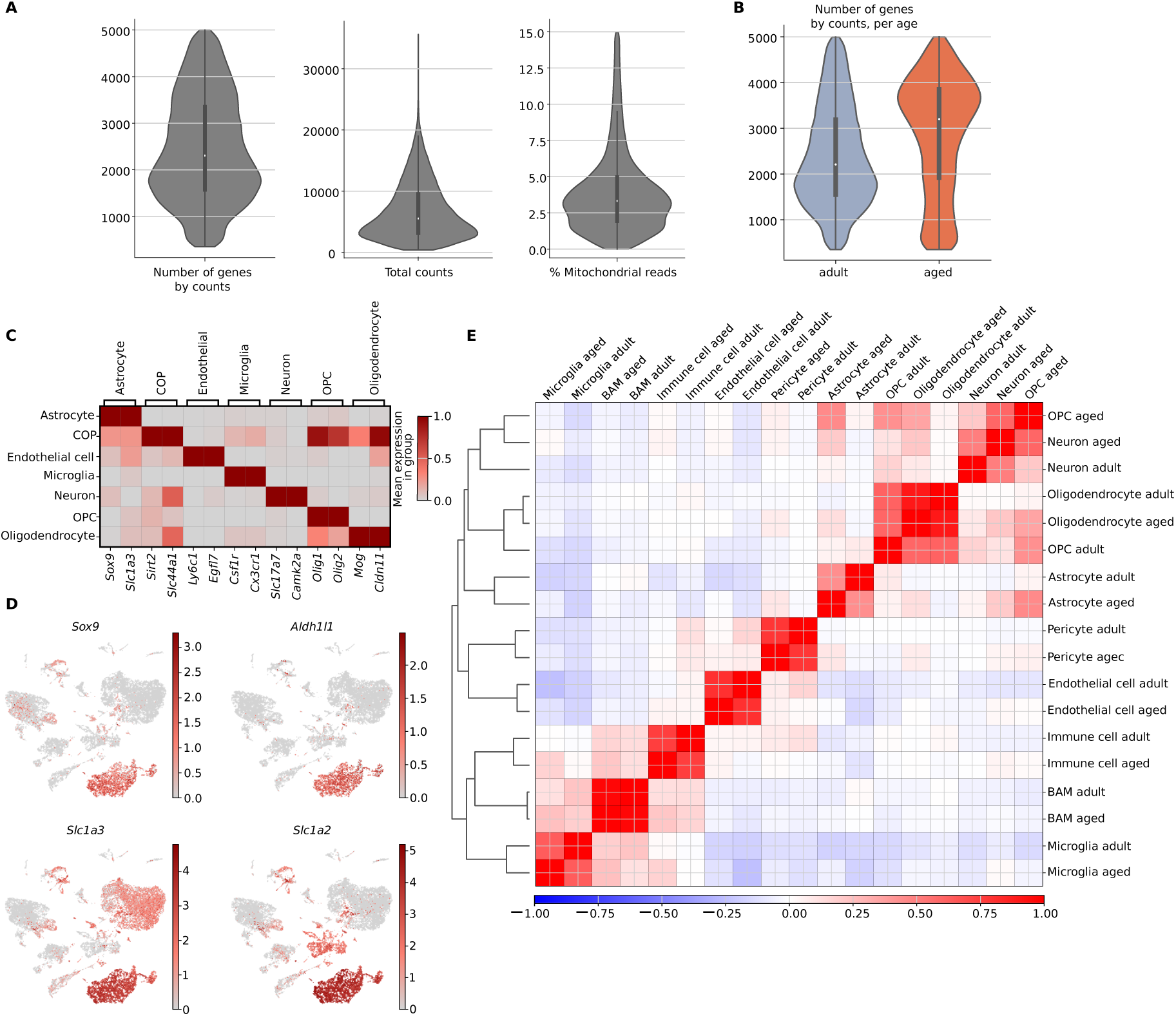
Single cell analysis of adult and aged cortical astrocytes. **(A)-(B)** Quality control parameters across the scRNA-seq dataset (A) and per condition (B). **(C)** Gene expression profiles of cell type-specific markers used for cluster annotation. BAM = border-associated macrophage, OPC = oligodendrocyte precursor cell. **(D)** Umap of common astrocyte markers. **(E)** Gene expression profile correlation matrix of all adult and aged cell types.

**Extended Data Figure 2.**
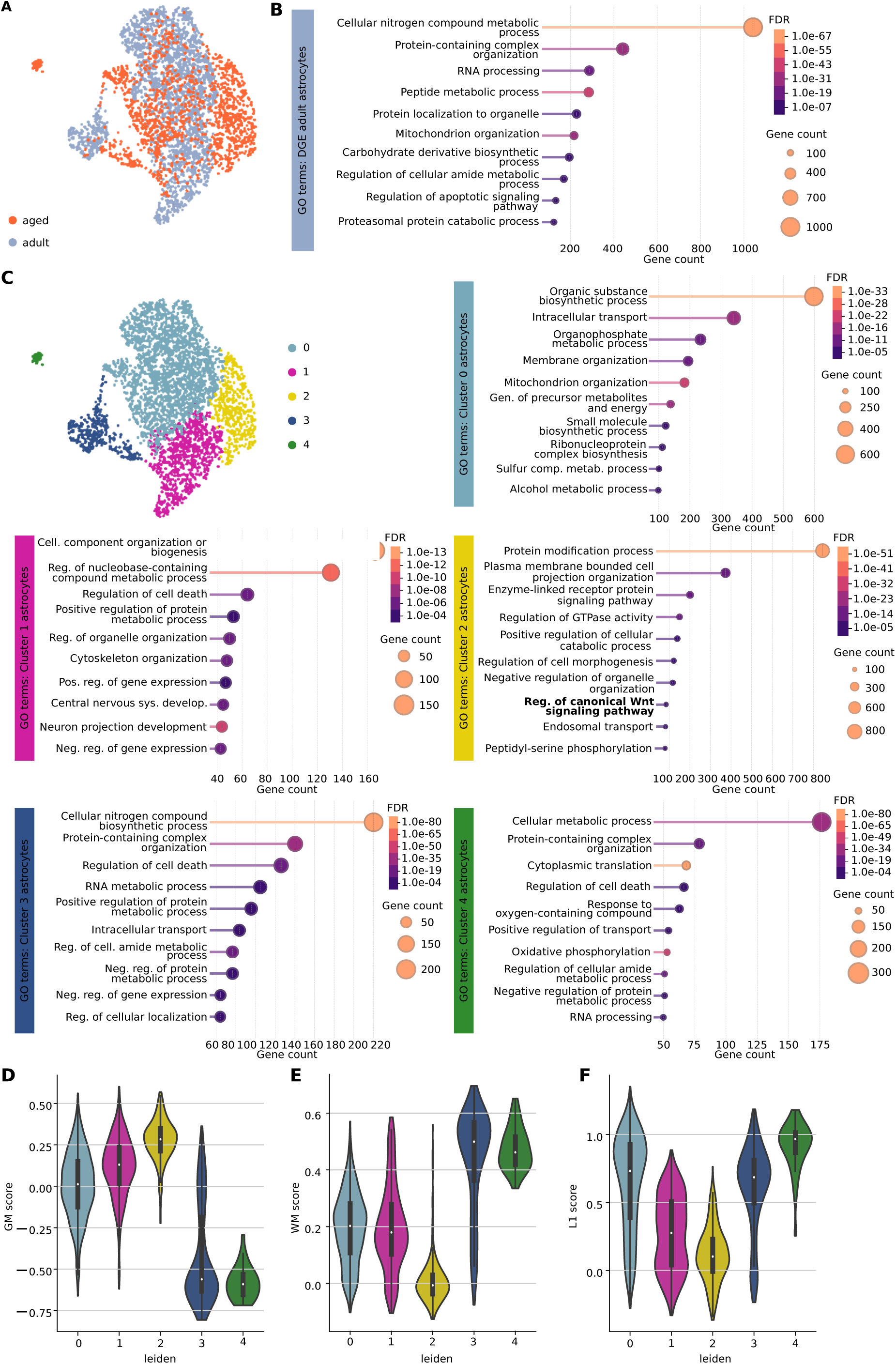
GO term analysis of the scRNA-seq. **(A)** Umap of astrocyte clusters from adult (color-coded in grey) and aged (color-coded in orange) astrocytes. **(B)** STRING-based GO terms of genes significantly enriched in adult compared to aged astrocytes. **(C)** Umap of 5 distinct color-coded astrocyte clusters and STRING-based GO terms of genes enriched in astrocyte clusters 0,1,2,3 and 4. **(D)–(F)** Violin plots of GM gene score (D), WM gene score (E) and L1 gene score (F) across astrocytes clusters.

**Extended Data Figure 3.**
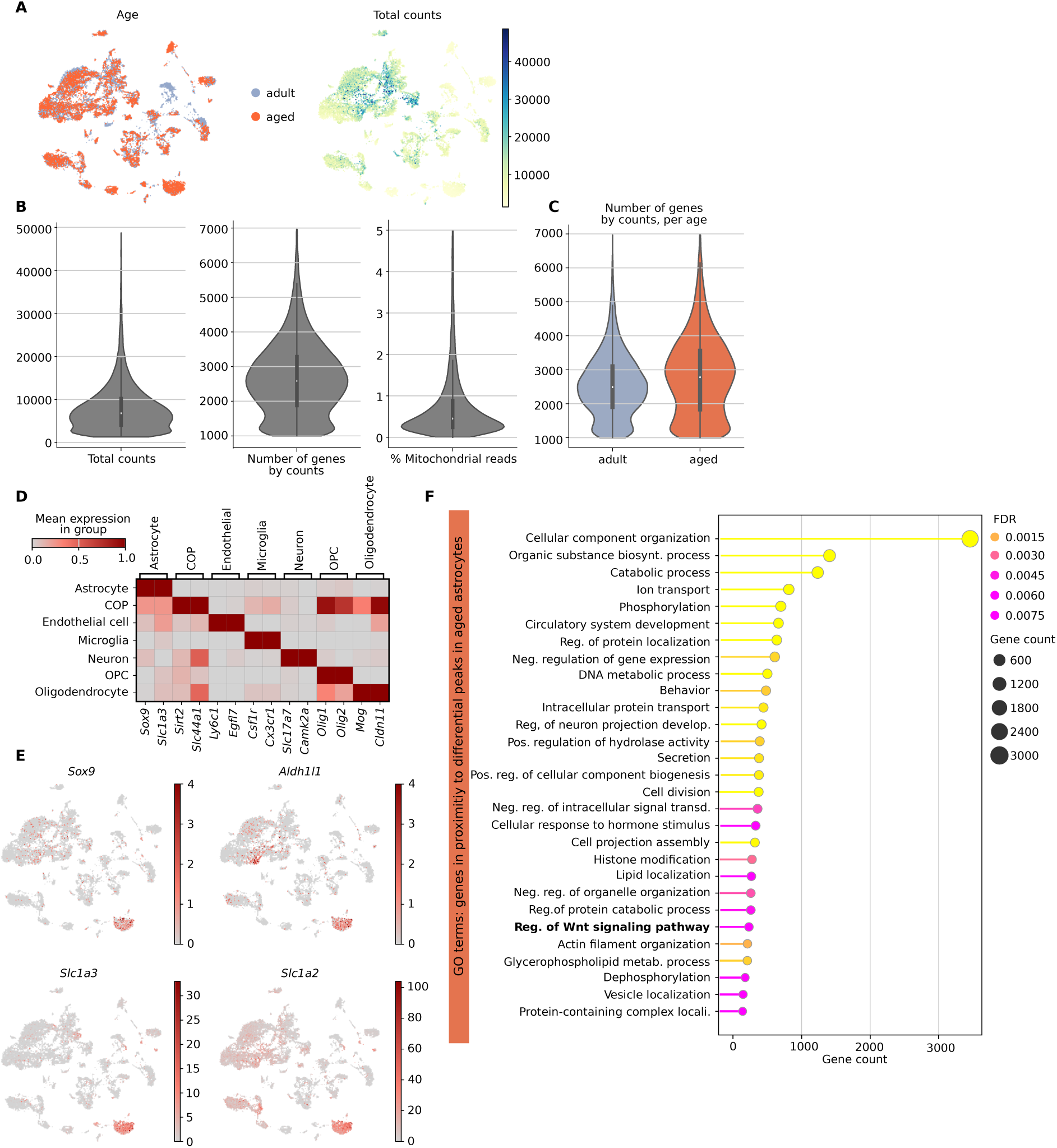
Analysis of the multiome dataset. **(A)** Umap representation of total RNA counts present in adult (color-coded in grey) and aged (color-coded in orange) from the multiome dataset. **(B)-(C)** Quality control parameters across the multiome RNA dataset (B) and per condition (C). **(D)** Matrix plot of cell type-specific markers used for cluster annotation. COP = committed oligodendrocyte precursor, OPC = oligodendrocyte precursor cell. **(E)** Umap of common astrocyte markers. **(F)** Extended list of STRING-based GO terms of genes associated to open regions in aged astrocytes.

**Extended Data Figure 4.**
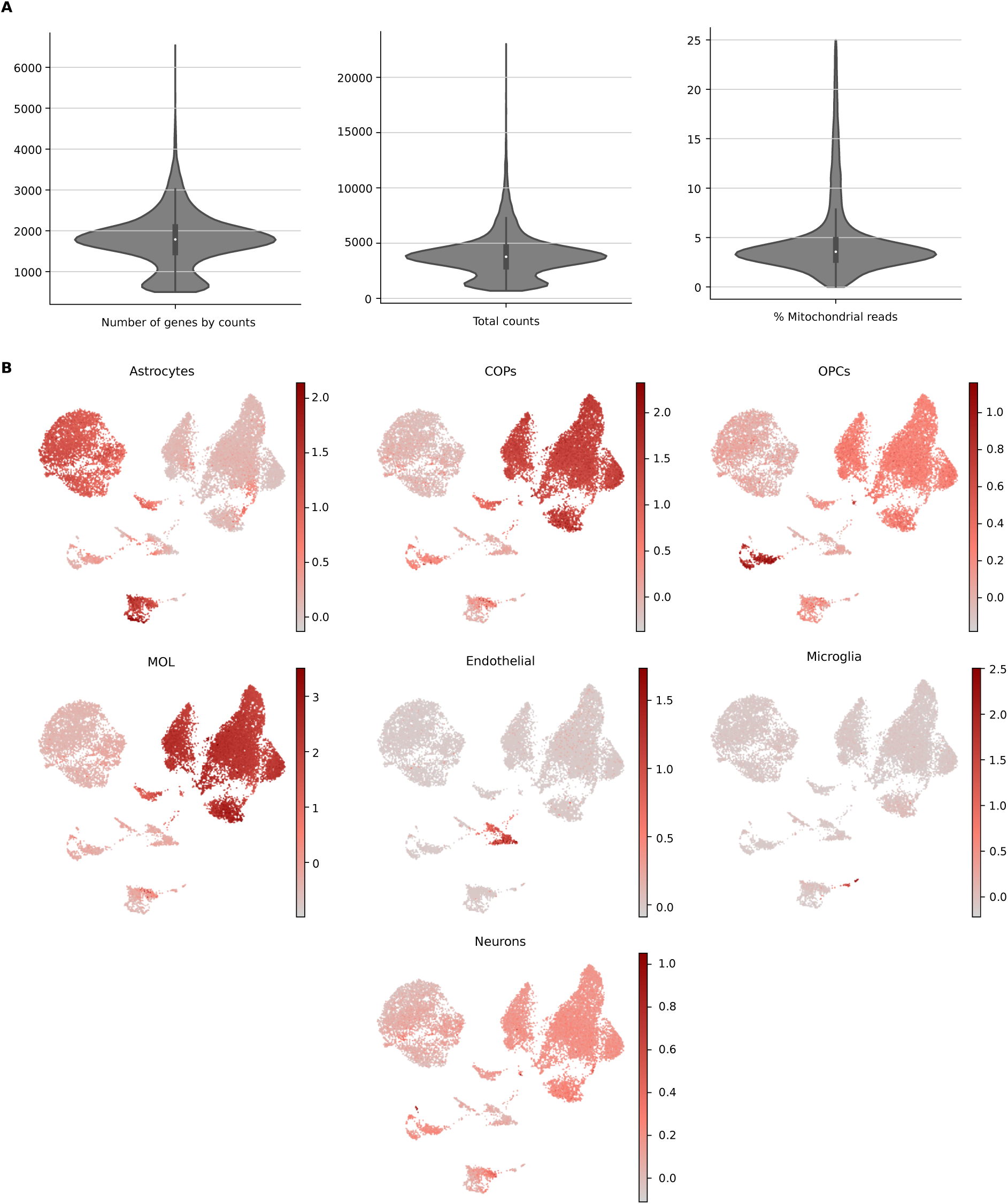
Quality parameters and cell type scores for the scRNA-seq dataset from cortical layers. **(A)** Violin plots of quality control parameters of the scRNA-seq dataset of cells isolated from different cortical layers. **(B)** Umaps showing marker gene scores for different cell types. COP = committed oligodendrocyte precursor, OPC = oligodendrocyte precursor cell, MOL = myelinating oligodendrocytes.

**Extended Data Figure 5.**
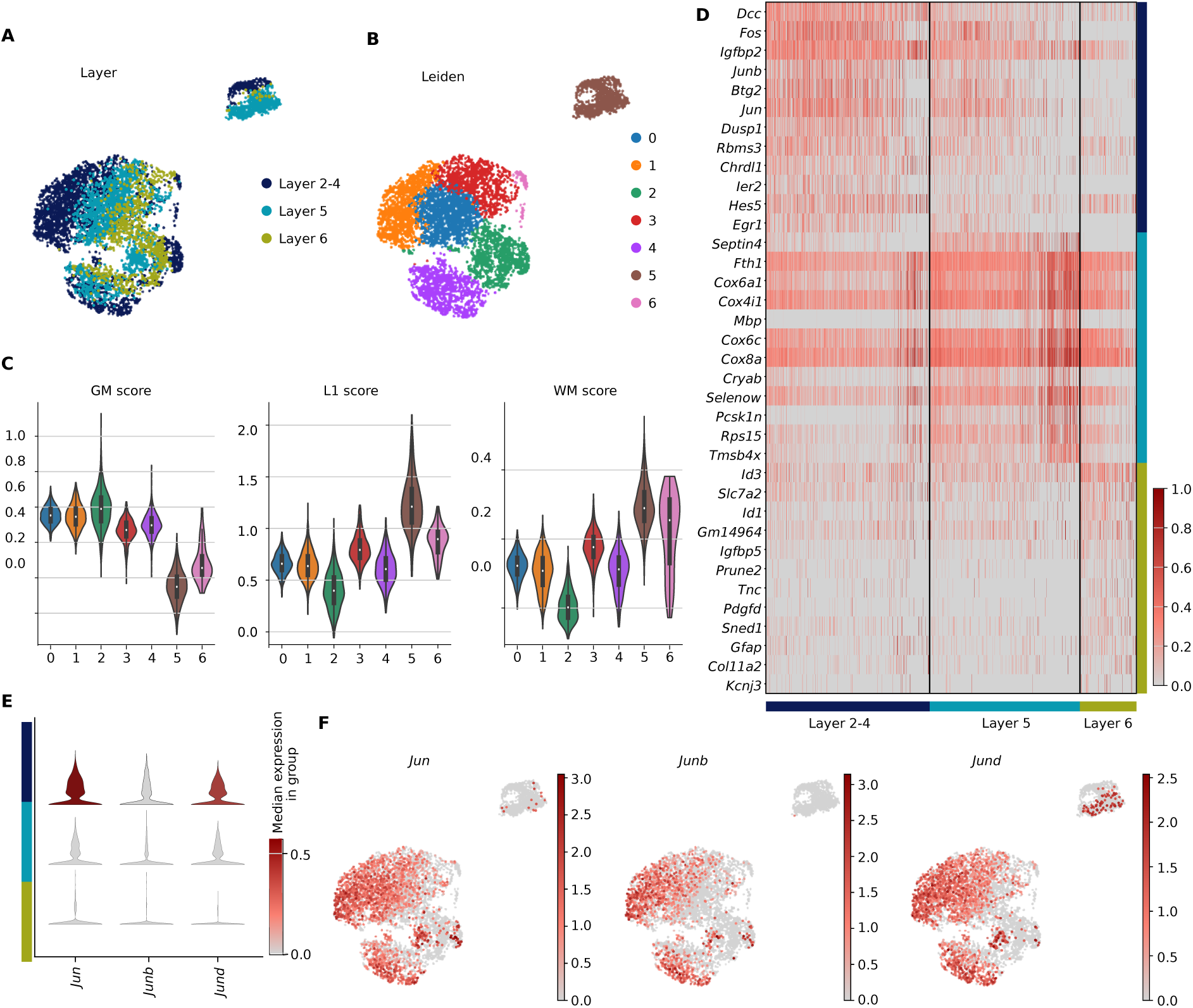
Layer dependent expression of Jun transcription factors. **(A)-(B)** Umap of scRNA-seq of astrocytes isolated from different cortical layers colored by layer **(A)** and unsupervised Leiden clustering (B). **(C)** Violin plots of GM, L1 and WM gene scores across astrocyte clusters. **(D)** Heatmap depicting the top 12 up-regulated genes per layer (Benjamini-Hochberg adjusted p-value < 0.05, unique per layer, expressed in at least 5 cells). **(E)** Violin plot showing the expression of *Jun*, *JunB* and *JunD* in astrocytes separated by layer. **(F)** Umaps showing expression of *Jun*, *JunB* and *JunD* in astrocytes.

## References

Allen, W.E. et al. (2023) ‘Molecular and spatial signatures of mouse brain aging at single-cell resolution’, Cell, 186(1), pp. 194–208.e18. Available at: 10.1016/j.cell.2022.12.010.

Arredondo, S.B. et al. (2020) ‘Role of Wnt Signaling in Adult Hippocampal Neurogenesis in Health and Disease’, Frontiers in Cell and Developmental Biology, Volume 8-2020. Available at: https://www.frontiersin.org/journals/cell-and-developmental-biology/articles/10.3389/fcell.2020.00860.

Bankhead, P. et al. (2017) ‘QuPath: Open source software for digital pathology image analysis’, Scientific Reports, 7(1), p. 16878. Available at: 10.1038/s41598-017-17204-5.

Barres, B.A. (2008) ‘The Mystery and Magic of Glia: A Perspective on Their Roles in Health and Disease’, Neuron, 60(3), pp. 430–440. Available at: 10.1016/j.neuron.2008.10.013.

Bengoa-Vergniory, N. et al. (2014) ‘A Switch From Canonical to Noncanonical Wnt Signaling Mediates Early Differentiation of Human Neural Stem Cells’, Stem Cells, 32(12), pp. 3196–3208. Available at: 10.1002/stem.1807.

Bocchi, R. et al. (2025) ‘Astrocyte heterogeneity reveals region-specific astrogenesis in the white matter’, Nature Neuroscience, 28(3), pp. 457–469. Available at: 10.1038/s41593-025-01878-6.

Boisvert, M.M. et al. (2018) ‘The Aging Astrocyte Transcriptome from Multiple Regions of the Mouse Brain’, Cell Reports, 22(1), pp. 269–285. Available at: 10.1016/j.celrep.2017.12.039.

Chien, J.-F. et al. (2024) ‘Cell-type-specific effects of age and sex on human cortical neurons’, Neuron, 112(15), pp. 2524–2539.e5. Available at: 10.1016/j.neuron.2024.05.013.

Chung, W.-S. et al. (2013) ‘Astrocytes mediate synapse elimination through MEGF10 and MERTK pathways’, Nature. 2013/11/24 edn, 504(7480), pp. 394–400. Available at: 10.1038/nature12776.

Clarke, L.E. et al. (2018) ‘Normal aging induces A1-like astrocyte reactivity’, Proceedings of the National Academy of Sciences, 115(8), p. E1896 LP–E1905. Available at: 10.1073/pnas.1800165115.

Danese, A. et al. (2021) ‘EpiScanpy: integrated single-cell epigenomic analysis’, Nature Communications, 12(1), p. 5228. Available at: 10.1038/s41467-021-25131-3.

Faust, T.E. et al. (2025) ‘Microglia-astrocyte crosstalk regulates synapse remodeling via Wnt signaling’, Cell, 188(19), pp. 5212–5230.e21. Available at: 10.1016/j.cell.2025.08.023.

Flores, N.M., et al. (2024) ‘Downregulation of Dickkopf-3, a Wnt antagonist elevated in Alzheimer’s disease, restores synapse integrity and memory in a disease mouse model’, eLife. Edited by B. Stevens and S.B. Nelson, 12, p. RP89453. Available at: 10.7554/eLife.89453.

Fu, H. et al. (2025) ‘Global burden of Alzheimer’s disease and other dementias (1990–2021): inequality, frontier, and decomposition analysis’, Frontiers in Aging Neuroscience, Volume 17-2025. Available at: https://www.frontiersin.org/journals/aging-neuroscience/articles/10.3389/fnagi.2025.1637029.

Gaikwad, S. et al. (2021) ‘Tau oligomer induced HMGB1 release contributes to cellular senescence and neuropathology linked to Alzheimer’s disease and frontotemporal dementia’, Cell Reports, 36(3). Available at: 10.1016/j.celrep.2021.109419.

González-Blas, C.B. et al. (2023) ‘SCENIC+: single-cell multiomic inference of enhancers and gene regulatory networks’, Nature Methods, 20(9), pp. 1355–1367. Available at: 10.1038/s41592-023-01938-4.

Gorelov, R. and Hochedlinger, K. (2024) ‘A cellular identity crisis? Plasticity changes during aging and rejuvenation’, Genes & Development [Preprint]. Available at: 10.1101/gad.351728.124.

Guérit, S. et al. (2021) ‘Astrocyte-derived Wnt growth factors are required for endothelial blood-brain barrier maintenance’, Progress in Neurobiology, 199, p. 101937. Available at: 10.1016/j.pneurobio.2020.101937.

Gujral, T.S. and MacBeath, G. (2010) ‘A System-Wide Investigation of the Dynamics of Wnt Signaling Reveals Novel Phases of Transcriptional Regulation’, PLOS ONE, 5(4), p. e10024. Available at: 10.1371/journal.pone.0010024.

Hahn, O. et al. (2023) ‘Atlas of the aging mouse brain reveals white matter as vulnerable foci’, Cell, 186(19), pp. 4117–4133.e22. Available at: 10.1016/j.cell.2023.07.027.

Heimann, G. et al. (2017) ‘Changes in the Proliferative Program Limit Astrocyte Homeostasis in the Aged Post-Traumatic Murine Cerebral Cortex’, Cerebral Cortex, 27(8), pp. 4213–4228. Available at: 10.1093/cercor/bhx112.

Heintz, N. (2004) ‘Gene Expression Nervous System Atlas (GENSAT)’, Nature Neuroscience, 7(5), p. 483. Available at: 10.1038/nn0504-483.

Hennes, M. et al. (2025) ‘Astrocyte diversity and subtypes: aligning transcriptomics with multimodal perspectives’, EMBO reports, 26(17), pp. 4203–4218–4218. Available at: 10.1038/s44319-025-00529-y.

Heo, D. et al. (2025) ‘Transcriptional profiles of mouse oligodendrocyte precursor cells across the lifespan’, Nature Aging, 5(4), pp. 675–690. Available at: 10.1038/s43587-025-00840-2.

Hwang, S.-G. et al. (2005) ‘Wnt-3a regulates chondrocyte differentiation via c-Jun/AP-1 pathway’, FEBS letters, 579(21), pp. 4837–4842. Available at: 10.1016/j.febslet.2005.07.067.

Inestrosa, N.C. et al. (2020) ‘Wnt Signaling Pathway Dysregulation in the Aging Brain: Lessons From the Octodon degus’, Frontiers in Cell and Developmental Biology, Volume 8- 2020. Available at: https://www.frontiersin.org/journals/cell-and-developmental-biology/articles/10.3389/fcell.2020.00734.

Jin, K. et al. (2025) ‘Brain-wide cell-type-specific transcriptomic signatures of healthy ageing in mice’, Nature, 638(8049), pp. 182–196. Available at: 10.1038/s41586-024-08350-8.

Jo, J. et al. (2021) ‘Regional heterogeneity of astrocyte morphogenesis dictated by the formin protein, Daam2, modifies circuit function’, The EMBO Reports, 22(12), p. EMBR202153200. Available at: 10.15252/embr.202153200.

Komkleow, S. et al. (2026) ‘Box A of HMGB1 plasmid reverses the age-related changes in the plasma proteomic profile of perimenopausal monkeys’, Scientific Reports [Preprint]. Available at: 10.1038/s41598-026-46747-9.

L. Lun, A.T., Bach, K. and Marioni, J.C. (2016) ‘Pooling across cells to normalize single-cell RNA sequencing data with many zero counts’, Genome Biology, 17(1), p. 75. Available at: 10.1186/s13059-016-0947-7.

Labarta-Bajo, L. and Allen, N.J. (2025) ‘Astrocytes in aging’, Neuron, 113(1), pp. 109–126. Available at: 10.1016/j.neuron.2024.12.010.

Lanjakornsiripan, D. et al. (2018) ‘Layer-specific morphological and molecular differences in neocortical astrocytes and their dependence on neuronal layers’, Nature Communications, 9(1), p. 1623. Available at: 10.1038/s41467-018-03940-3.

Lee, E. et al. (2022) ‘A distinct astrocyte subtype in the aging mouse brain characterized by impaired protein homeostasis’, Nature Aging, 2(8), pp. 726–741. Available at: 10.1038/s43587-022-00257-1.

Ling, E. et al. (2024) ‘A concerted neuron–astrocyte program declines in ageing and schizophrenia’, Nature, 627(8004), pp. 604–611. Available at: 10.1038/s41586-024-07109-5.

Linker, K.E. et al. (2025) ‘Aging in mice alters regionally enriched striatal astrocytes’, Nature Communications, 16(1), p. 8496. Available at: 10.1038/s41467-025-63429-8.

Liu, H. et al. (2022) ‘Molecular and cytological profiling of biological aging of mouse cochlear inner and outer hair cells’, Cell Reports, 39(2). Available at: 10.1016/j.celrep.2022.110665.

Liu, J. et al. (2022) ‘Wnt/β-catenin signalling: function, biological mechanisms, and therapeutic opportunities’, Signal Transduction and Targeted Therapy, 7(1), p. 3. Available at: 10.1038/s41392-021-00762-6.

Liu, K. et al. (2024) ‘The decreased astrocyte-microglia interaction reflects the early characteristics of Alzheimer’s disease’, iScience, 27(3). Available at: 10.1016/j.isci.2024.109281.

Liu, W.-S. et al. (2025) ‘Plasma proteomics identify biomarkers and undulating changes of brain aging’, Nature Aging, 5(1), pp. 99–112. Available at: 10.1038/s43587-024-00753-6.

López-Otín, C. et al. (2023) ‘Hallmarks of aging: An expanding universe’, Cell, 186(2), pp. 243–278. Available at: 10.1016/j.cell.2022.11.001.

Matho, K.S. et al. (2021) ‘Genetic dissection of the glutamatergic neuron system in cerebral cortex’, Nature, 598(7879), pp. 182–187. Available at: 10.1038/s41586-021-03955-9.

McBean, G.J. (2018) ‘Astrocyte Antioxidant Systems’, Antioxidants. Available at: 10.3390/antiox7090112.

Mertens, J. et al. (2021) ‘Age-dependent instability of mature neuronal fate in induced neurons from Alzheimer’s patients’, Cell Stem Cell, 28(9), pp. 1533–1548.e6. Available at: 10.1016/j.stem.2021.04.004.

Morrison, J.H. and Baxter, M.G. (2012) ‘The ageing cortical synapse: hallmarks and implications for cognitive decline’, Nature Reviews Neuroscience, 13(4), pp. 240–250. Available at: 10.1038/nrn3200.

Natarajan, P. et al. (2024) ‘Single Cell Deletion of the Transcription Factors Trps1 and Sox9 in Astrocytes Reveals Novel Functions in the Adult Cerebral Cortex’, Glia, n/a(n/a). Available at: 10.1002/glia.24645.

Ogamino, S. et al. (2024) ‘Dynamics of Wnt/β-catenin reporter activity throughout whole life in a naturally short-lived vertebrate’, npj Aging, 10(1), p. 23. Available at: 10.1038/s41514-024-00149-1.

Ohlig, S. et al. (2021) ‘Molecular diversity of diencephalic astrocytes reveals adult astrogenesis regulated by Smad4’, The EMBO Journal, 40(21), p. e107532. Available at: 10.15252/embj.2020107532.

Organization, W.H. (2021) ‘Ageing and health’. Available at: https://www.who.int/news-room/fact-sheets/detail/ageing-and-health.

Paneni, F. et al. (2013) ‘Deletion of the Activated Protein-1 Transcription Factor JunD Induces Oxidative Stress and Accelerates Age-Related Endothelial Dysfunction’, Circulation, 127(11), pp. 1229–1240. Available at: 10.1161/CIRCULATIONAHA.112.000826.

Park, S.Y. et al. (2020) ‘SPON1 Can Reduce Amyloid Beta and Reverse Cognitive Impairment and Memory Dysfunction in Alzheimer’s Disease Mouse Model’, Cells, 9(5). Available at: 10.3390/cells9051275.

Peng, S. et al. (2025) ‘Global, regional and national burden of Parkinson’s disease in people over 55 years of age: a systematic analysis of the global burden of disease study, 1991– 2021’, BMC Neurology, 25(1), p. 178. Available at: 10.1186/s12883-025-04191-8.

Popov, A. et al. (2021) ‘Astrocyte dystrophy in ageing brain parallels impaired synaptic plasticity’, Aging Cell, 20(3), p. e13334. Available at: 10.1111/acel.13334.

Popov, A. et al. (2023) ‘Mitochondrial malfunction and atrophy of astrocytes in the aged human cerebral cortex’, Nature Communications, 14(1), p. 8380. Available at: 10.1038/s41467-023-44192-0.

Saadeddin, A. et al. (2009) ‘The links between transcription, beta-catenin/JNK signaling, and carcinogenesis’, Molecular cancer research: MCR, 7(8), pp. 1189–1196. Available at: 10.1158/1541-7786.MCR-09-0027.

Allen, N.J. (2014) ‘Astrocyte Regulation of Synaptic Behavior’, Annual Review of Cell and Developmental Biology, 30(1), pp. 439–463. Available at: 10.1146/annurev-cellbio-100913-013053.

Salminen, A. (2026) ‘Redox-sensitive high mobility group box 1 (HMGB1) protein is a multipotent regulator in the pathogenesis of Alzheimer’s disease’, Neurochemistry International, 193, p. 106115. Available at: 10.1016/j.neuint.2026.106115.

Shaulian, E. and Karin, M. (2002) ‘AP-1 as a regulator of cell life and death’, Nature Cell Biology, 4(5), pp. E131–E136. Available at: 10.1038/ncb0502-e131.

Siletti, K. et al. (2025) ‘Transcriptomic diversity of cell types across the adult human brain’, Science, 382(6667), p. eadd7046. Available at: 10.1126/science.add7046.

Sinha, A. et al. (2025) ‘Repression of Wnt/β-catenin signaling by SOX9 and Mastermind-like transcriptional coactivator 2’, Science Advances, 7(8), p. eabe0849. Available at: 10.1126/sciadv.abe0849.

Sofiadis, K. et al. (2021) ‘HMGB1 coordinates SASP-related chromatin folding and RNA homeostasis on the path to senescence’, Molecular Systems Biology, 17(6), p. e9760. Available at: 10.15252/msb.20209760.

Solovey, M. et al. (2024) ‘Community assesses differential cell communication using large multi-sample case-control scRNAseq datasets’, bioRxiv, p. 2024.03.01.582941. Available at: 10.1101/2024.03.01.582941.

Stogsdill, J.A. et al. (2022) ‘Pyramidal neuron subtype diversity governs microglia states in the neocortex’, Nature, 608(7924), pp. 750–756. Available at: 10.1038/s41586-022-05056-7.

Supasai, S. et al. (2026) ‘HMGB1 Box A gene therapy reverses cognitive and neuropathological features in AlCl₃/D-galactose rat model of Alzheimer’s disease’, Experimental Neurology, 397, p. 115583. Available at: 10.1016/j.expneurol.2025.115583.

Tartiere, A.G., Freije, J.M.P. and López-Otín, C. (2024) ‘The hallmarks of aging as a conceptual framework for health and longevity research’, Frontiers in Aging, Volume 5-2024. Available at: https://www.frontiersin.org/journals/aging/articles/10.3389/fragi.2024.1334261.

Thrupp, N. et al. (2020) ‘Single-Nucleus RNA-Seq Is Not Suitable for Detection of Microglial Activation Genes in Humans’, Cell Reports, 32(13). Available at: 10.1016/j.celrep.2020.108189.

Toualbi, K. et al. (2007) ‘Physical and functional cooperation between AP-1 and β-catenin for the regulation of TCF-dependent genes’, Oncogene, 26(24), pp. 3492–3502. Available at: 10.1038/sj.onc.1210133.

Vainchtein, I.D. et al. (2018) ‘Astrocyte-derived interleukin-33 promotes microglial synapse engulfment and neural circuit development’, Science, 359(6381), pp. 1269–1273. Available at: 10.1126/science.aal3589.

Verkhratsky, A. et al. (2012) ‘Neurological diseases as primary gliopathies: a reassessment of neurocentrism’, ASN neuro, 4(3), p. e00082. Available at: 10.1042/AN20120010.

Verkhratsky, A., et al. (2021) ‘Astrocytes: The Housekeepers and Guardians of the CNS BT - Astrocytes in Psychiatric Disorders’, in B. Li et al. (eds). Springer International Publishing, pp. 21–53. Available at: 10.1007/978-3-030-77375-5_2.

Verkhratsky, A. et al. (2023) ‘Astrocytes in human central nervous system diseases: a frontier for new therapies’, Signal Transduction and Targeted Therapy, 8(1), p. 396. Available at: 10.1038/s41392-023-01628-9.

Wang, C.-Y. et al. (2023) ‘Daam2 phosphorylation by CK2α negatively regulates Wnt activity during white matter development and injury’, Proceedings of the National Academy of Sciences, 120(35), p. e2304112120. Available at: 10.1073/pnas.2304112120.

Weitzman, J.B. et al. (2000) ‘JunD Protects Cells from p53-Dependent Senescence and Apoptosis’, Molecular Cell, 6(5), pp. 1109–1119. Available at: 10.1016/S1097-2765(00)00109-X.

Wolf, F.A., Angerer, P. and Theis, F.J. (2018) ‘SCANPY: large-scale single-cell gene expression data analysis’, Genome Biology, 19(1), p. 15. Available at: 10.1186/s13059-017-1382-0.

Wu, C. et al. (2024) ‘Spatially resolved transcriptome of the aging mouse brain’, Aging Cell, 23(5), p. e14109. Available at: 10.1111/acel.14109.

Ximerakis, M. et al. (2019) ‘Single-cell transcriptomic profiling of the aging mouse brain’, Nature Neuroscience, 22(10), pp. 1696–1708. Available at: 10.1038/s41593-019-0491-3.

Xue, C. et al. (2025) ‘Wnt signaling pathways in biology and disease: mechanisms and therapeutic advances’, Signal Transduction and Targeted Therapy, 10(1), p. 106. Available at: 10.1038/s41392-025-02142-w.

Yang, J.-H. et al. (2023) ‘Loss of epigenetic information as a cause of mammalian aging’, Cell, 186(2), pp. 305–326.e27. Available at: 10.1016/j.cell.2022.12.027.

Zhang, L. et al. (2025) ‘Modulating mTOR-dependent astrocyte substate transitions to alleviate neurodegeneration’, Nature Aging [Preprint]. Available at: 10.1038/s43587-024-00792-z.

